# Hair phenotype diversity across Indriidae lemurs

**DOI:** 10.1101/2021.10.16.464615

**Authors:** Elizabeth Tapanes, Rachel L. Jacobs, Ian Harryman, Edward E. Louis, Mitchell T. Irwin, Jason M. Kamilar, Brenda J. Bradley

## Abstract

**Objectives:** Hair (i.e., pelage/fur) is a salient feature of primate (including human) diversity and evolution— serving functions tied to thermoregulation, protection, camouflage, and signaling—but wild primate pelage evolution remains relatively understudied. Specifically, assessing multiple hypotheses across distinct phylogenetic scales is essential but is rarely conducted. We examine whole body hair color and density variation across Indriidae (*Avahi*, *Indri*, *Propithecus*)—a lineage that, like humans, exhibits vertical posture (i.e., their whole bodies are vertical to the sun).

**Materials and Methods:** Our analyses consider multiple phylogenetic scales (family-level, genus-level) and hypotheses (e.g., Gloger’s rule, the body cooling hypotheses). We obtain hair color and density from museum and/or wild animals, opsin genotypes from wild animals, and climate data from WorldClim. To analyze our data, we use phylogenetic generalized linear mixed models (PGLMM) using Markov chain Monte Carlo algorithms.

**Results:** Our results show that across the Indriidae family, darker hair is typical in wetter regions. However, within *Propithecus*, dark black hair is common in colder forest regions. Results also show pelage redness increases in populations exhibiting enhanced color vision. Lastly, we find follicle density on the crown and limbs increases in dry and open environments.

**Discussion:** This study highlights how different selective pressures across distinct phylogenetic scales have likely acted on primate hair evolution. Specifically, our data across *Propithecus* may implicate thermoregulation and is the first empirical evidence of Bogert’s rule in mammals. Our study also provides rare empirical evidence supporting an early hypothesis on hominin hair evolution.

## Introduction

Hair (i.e., pelage, fur) is a salient feature of primate diversity and evolution and is the product of natural and sexual selection. Hair has been found to play a role in conspecific signaling (Allen & Higham, 2013), thermoregulation (Fratto & Davis, 2011), camouflage (Barrett et al., 2019; Hoekstra et al., 2005) and to possibly provide protection from pathogens/parasites (Paus & Foitzik, 2004). Across all mammals, primates—including humans— arguably exhibit the most diversity in hair color (e.g., bright reds) and growth patterns (e.g., mustaches) between and within species (Caro, 2005; Rowe, 1996). For example, within Colobinae, some *Colobus* spp. sport a long black and white dorsal cape, and *Pygathrix* spp. are often tri-colored and short-haired (Rowe, 1996). Similarly, human hair color and texture vary throughout the human body, between individuals, between populations, and can be correlated with ancestry and ethnicity (Lasisi et al., 2016; Seibert & Steggerda, 1999; Steggerda & Seibert, 1941). Nevertheless, our knowledge about hair evolution is based mainly on non-primate mammals. Primate hair remains relatively understudied (with some notable exceptions: (e.g., (Allen & Higham, 2013; Bell et al., 2021; Kamilar et al., 2013; Rakotonirina et al., 2017; Santana et al., 2012)), with most research taking a macro-scale approach and sampling broadly across the primate order (e.g., all catarrhines).

These previous studies indicate that variation in primate hair color likely evolved in response to multiple distinct but not mutually exclusive pressures associated with ecology, visual systems, activity cycles, and sociality (Allen & Higham, 2013; Bell et al., 2021; Kamilar et al., 2013; Rakotonirina et al., 2017; Santana et al., 2012). For example, across primates, melanic (i.e., black) colors often co-occur with humid regions—which may be due to camouflage, pleiotropy, pathogen resistance, or selection on a related trait (i.e., Gloger’s rule) (Bell et al., 2021; Kamilar & Bradley, 2011; Santana et al., 2012; 2013). Gloger’s rule generally predicts birds and mammals will be more pigmented in tropical regions where it is warm and humid, such as tropical rainforests (Gloger, 1833). However, without accounting for variation within families, genera, and species, other subtle trends or relationships may not be detected. For example, recent work in birds suggests a distinct rule (Bogert’s rule) may explain some melanic variation in endotherms depending on the phylogenetic and geographic scale sampled (Bogert, 1949; Delhey et al., 2019). According to Bogert’s rule, melanic colors should co-occur with cold regions as dark colors may absorb heat better and aid in thermoregulation (Bogert, 1949; Delhey et al., 2019). However, there is no current empirical evidence for this rule in primates (or any mammals). Yet across human populations, variation in eumelanin exceeds that seen in red, blonde, or lighter hair (Lasisi et al., 2016), which suggests that important biological variation may be identified if the scale of analyses was smaller than previously considered (i.e., within a few populations vs. across all populations) (Norton et al., 2016).

Across non-human primates, red hues (and facial patterning) may act as a means to aid conspecific communication (Cuthill et al., 2017; Santana et al., 2012; 2013; Winters et al., 2020). For example, diurnal activity and the ability to perceive a wider range of colors compared to other mammals may have contributed to the enhanced color diversity in specific primate clades (Bell et al., 2021). However, the evidence for coevolution of hair coloration, specifically red hair and color vision in primates, remains mixed (Fernandez & Morris, 2007; Kamilar et al., 2013). Our understanding of primate hair color may benefit from broader sampling across wild populations and/or analyses conducted on smaller phylogenetic scales.

Similarly, our understanding of pelage density evolution remains limited to extensive macro-scale studies. Across primates, the primary trend is that hair density decreases as body size increases (Sandel, 2013; Schwartz & Rosenblum, 1981). One early hypothesis argued that a reduction in body hair density across hominins coupled to increases in activity level, sweat glands, sweat production, and a change to a dry savannah environment may have aided thermoregulation (i.e., “body-cooling hypothesis”) (Jablonski & Chaplin, 2000; Post et al., 1975; Wheeler, 1992). Despite various physiological models testing evidence of the body-cooling hypothesis (Chaplin et al., 1994; Schwartz & Rosenblum, 1981; Wheeler, 1992), there is only rare empirical evidence for this phenomenon within primates (Best et al., 2019). There also remains a poor understanding of how hair density in primates may be impacted by other factors, such as climate. Because hair does not preserve in the fossil record, studies on wild primates can provide critical insights into the evolutionary mechanisms underlying one of the most interesting traits among humans.

This study examined whole-body pelage color and hair/follicle density across Indriidae— a diverse clade of lemurs. Since the Indriidae clade varies in critical aspects known to drive primate (including human) hair variation, they make a good proxy for testing multiple hair hypotheses that have not yet been jointly assessed in one study. Our overarching goals were to 1) examine if Indriidae pelage variation varies across distinct phylogenetic scales and 2) determine if multiple, non-exclusive selective pressures can impact the same trait. Specifically, we aimed to jointly assess the impacts of climate, body size, and color vision on hair evolution at distinct phylogenetic scales. Collectively, this family (comprising three genera: nocturnal *Avahi* and diurnal *Indri* and *Propithecus*) inhabits almost every bio-climate of Madagascar (e.g., humid rainforest, temperate forests, open spiny forest, mangroves) (Mittermeier et al., 2010). The hair of the 13 species in our study varies intra- and interspecifically in both color (e.g., melanic, tricolored orange) and growth morphologies (e.g., long silky hairs, short hairs) (King et al., 2014; Mittermeier et al., 2010; Petter & Peyrieras, 1972; Rakotonirina et al., 2014; Tattersall, 1986). Indriidae also vary in body size (≈1kg (*Avahi*), ≈3-8kg (*Propithecus*), ≈10kg (*Indri*))—and phylogenetic relatedness likely impacts body size evolution (Lehman, 2007). Thus, we specifically (1a\2a) aimed to uncover how climate and body size influences: pelage brightness (i.e., black color), hue (i.e., red color), and hair follicle density across Indriidae and within *Propithecus*. We expected that climate impacts pelage color and density evolution distinctly across different phylogenetic scales. We predicted that body size only impacts hair/follicle density, based on evidence from other primates (Sandel, 2013). Diurnal Indriidae (*Indri* and *Propithecus*) also vary in their color vision. In many species, some females have trichromatic color vision (ability to distinguish reddish and greenish hues), while males and other females are red-green color blind (polymorphic color vision; Jacobs et al., 2017). Although this feature occurs in some other day-active lemurs, Indriidae has been found to exhibit greater genetic variation in color vision across species (Jacobs et al., 2017). Accordingly, we (2b) aimed to understand how this variation in color vision impacts pelage hue in diurnal genera. We predicted that populations with enhanced capacity for trichromatic color vision would have individuals with redder pelage (Sumner & Mollon, 2003). Our results provide critical insight into how multiple variables impact primate hair evolution across scales. Importantly, this family consists solely of vertical clingers and leapers, which means they exhibit upright locomotory posture similar to humans (via bipedal hopping (Schmidt, 2011)). Thus, these results (specifically those on hair density) may provide insight into human hair evolution.

## Materials and Methods

### Hair morphology and body size data collection

We used hair color and density measurements from museum research skins (N_Total_ = 63; N_Museum_ = 59) and living animals in one wild population (N_Wild_ = 4, *P. diadema* from Tsinjoarivo) (SI 1). Museum samples spanned three museum collections (SI 2). In total, we sampled six *Avahi* populations (Figure 1A), 23 *Propithecus* populations (Figure 1B), and four *Indri* populations (Figure 1C). Our sampling included nocturnal and diurnal genera (Figure 1D) across all significant bioclimatic regions in Madagascar (Figure 1E). Many individuals did not have available age or sex-class data. Hence, we did not test for differences between age classes or sex. Although some preliminary evidence suggests there may be subtle pelage dichromatism, sifakas are unlikely highly sexually dichromatic in terms of coloration. Instead, sifakas exhibit more significant visible variation within sexes (i.e., bimorphism in males) (Lewis & Van Schaik, 2007; Mittermeier et al., 2010; Spriggs, 2017; Tapanes et al., 2018). Measurements of hair from wild sifakas consisted of following the same protocol as museum individuals. We also extracted body length (mm), measured from head to tip of tail, from all individuals as a proxy for body size using Photoshop CC 6. We standardized all measurements using the known length of a ruler in the photos. Wild sifakas were captured for ongoing parallel research using a standard capture protocol established by the Prosimian Biomedical Survey Project (PBSP) Team (used on 750+ lemurs, 16+ sites, and 35+ species since 2000) (Dutton et al., 2003; Irwin et al., 2010). Groups were habituated before capture. The capture team was led by a trained professional, and captures were performed at close range using Pneu-dart 9mm disposable non-barbed darts containing tiletamine/zolazepam (Telazol) for immobilization.

**Figure 1.**
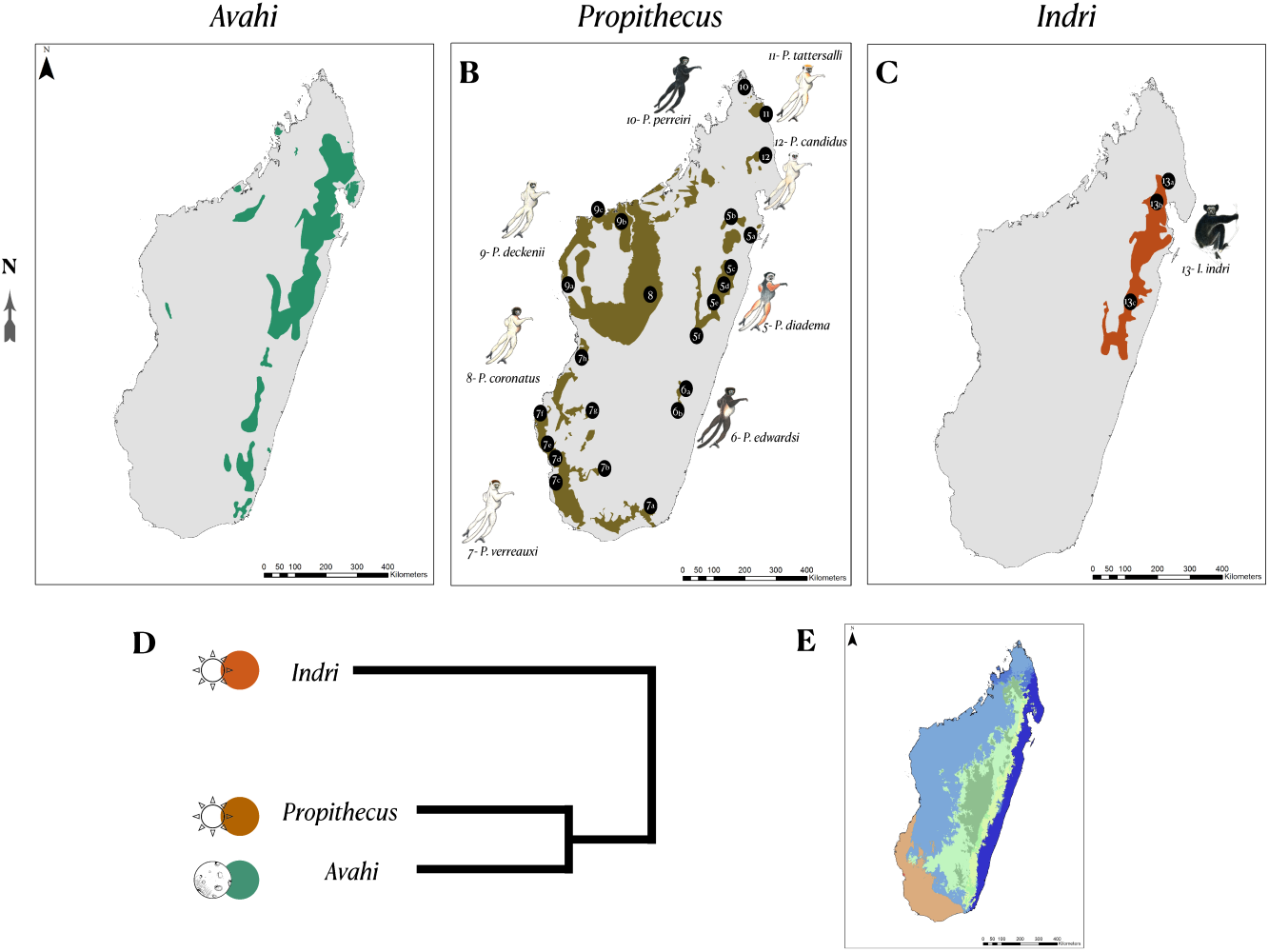
Indriidae species ranges and sampling localities for (A) *Avahi*, (B) *Propithecus*, and (C) *Indri*; along with a (D) cladogram, and (E) Koppen-climate zones that span the genera’s distribution. Key: *Avahi* spp. = (1a) Betampona, (1b) Sihanaka forest, (1c) Andasibe, (2) Forest west of Fort Dauphin, (3) Vondrozo, (4) Ankarafatsika; *Propithecus* spp. = (5a) Betampona, (5b) Zahamahena, (5c) Didy, (5d) Sihanaka forest, (5e) forest patch near Andasibe, (5f) Tsinjoarivo (eastern and western), (6a) Nandihizana, (6b) East of Farohny River, (7a) Mandrare Valley, (7b) Bevala East, (7c) Tsinanampetsotsa, (7d) Lower Ifaty, (7e) Ifaty, (7f) Lake Ihortry, (7g) Ankazoabo West, (7h) Upper Tsirbihina River, (8) Ambararatabe, (9a) Tsingy de Bemaraha, (9b) Tsingy de Namoraka, (9c) Baie de Baly Nat Park, (10) Reserve of Analamerana, (11) Forest fragment near Daraina, (12) Anjanaharibe-Sud Reserve; *Indri* indri = (13a) Anjanaharibe-Sud Reserve, (13b) 40 km West of Maroantsetra, (13c) Sihanaka forest

### Hair coloration collection

We used color-calibrated photographs of all animals (N = 63 individuals; SI 1) and a Canon EOS Rebel SL1/EOS 100D mounted with a Canon EF-S18-55mm f/3.5-5.6 IS STM lens. We calibrated each image against a color standard with 24 color squares of known reflectance values. The camera color mode was set to ’faithful.’ The camera was calibrated with a grey standard to set a manual white balance. All individuals were photographed against dull backgrounds using the adjacent method (Bergman & Beehner, 2008; Kendal et al., 2013). We followed established protocols for calibrating photos in previous studies for both museum and wild specimens (Stevens et al., 2007; 2009).

We used the Red-Green-Blue (RGB) color model and extracted all RGB values using Photoshop CC 6 across 14 body regions. Due to the high degree of variation across the pelage of the *Propithecus* tail, ventral surface, and dorsal surface, we divided measurements of these regions into three quadrants (lower, mid, and upper) (SI 3). If one of these regions was damaged, we took measurements from the center as the “average.” Primate hair occupies a relatively small color space, primarily varying along two axes. The first relates to variation in luminance and exhibits little chromatic variation and is captured by similar red, green, and blue values. Thus, we used RGB averages here as a proxy for ’brightness’ in a hair patch to indicate if the hair is white, a shade of grey, or black. The second axis of variation is hue and indicates variation along a red/brown axis and can be captured by a ratio of red to green. The latter is because the green and blue channels are most similar, while the red channel exhibits higher values on average. As the Red-Green ratio increases, hair appears increasingly red (Kamilar et al., 2013). We ran a correlation matrix across color values and removed any that were highly correlated with each other (r > .90). This reduced our hair color variables to RGB averages and RG ratios for the (1) crown, (2) cheek, (3) dorsal torso, (4) ventral torso, (5) forelimb, (6) hindlimb, and (7) tail. In total, we collected 882 brightness and 882 hue measurements (63 individuals x 14 sampling regions). Previous work on museum skins indicated no effect of storage time on pelage coloration from 142 museum skins collected over 100 years (1898 and 1998) (Kamilar et al., 2013).

### Hair and follicle density collection

We used a protocol that borrows from dermatological methods and instrumentation, which automates hair density measurements (Avram & Rogers, 2009). We used CompareViewHair® software (STR Technologies) tools combined with a specialized hand-held digital light microscope (ProScope HR2©; Bodelin) with a 100x lens. The software included a built-in procedure for calibrating the lens and software before use. The hair of an animal was parted, and the lens was placed on the skin. A light within the microscope illuminated the skin and provided needed light for subsequently photographing the area. We collected microscopy images (detailing transects of hair follicles and shafts) and compiled semi-automated measurements in the laboratory using this system. We examined two hair morphology properties: hair density (shafts/cm^2^) and follicle density (infundibula/cm^2^).

We collected three microscopy photos (replicates) from five body regions (SI 4) of each animal (n = 47 individuals) (SI 1) using the hand-held microscope, adhering to a previously established protocol (Bradley et al., 2014). The method provided a visual transect/plot of follicles (i.e., infundibula) and the number of hair shafts emerging from each. We used the ‘density tool’ in CompareViewHair® software to estimate the number of infundibula per cm^2^ (i.e., follicle density). We marked each follicle strand manually and used that number to estimate the final value in any given region (given the specific magnification). However, this protocol assumed that the entire photograph was visible, which was not always the case. Therefore, in our final measurements for ’follicle density,’ we used the value the software provided of density per cm^2^ and divided it by the visible photo’s proportion. Similarly, for our final measurement for ‘hair density,’ we used the value the software calculated for density per cm^2^ and multiplied it by the average number of hairs emerging from each follicle. In total, we collected 235 hair and 235 follicle density measurements (47 individuals x 5 body regions). Previous work on museum skins shows no effect of storage time on pelage density for skins collected over 100 years (Bradley et al., 2014).

### Climatic and phylogenetic data collection

We collected climate and environmental data from 33 sampling locations across Madagascar (Figure 1). Specifically, we extracted data from WorldClim (version 1) that consisted of averages from 1960 to 1990. This likely created some noise in the data given museum data spanned the early 1900s to late 1990s. However, Indriidae lifespans (≈20+ years) and generation times are long (≈3-10 years) (Lawler et al., 2009; Godfrey et al., 2001; Guevara et al. 2018). Thus, the data are likely representative of accurate averages across bioclimates and phenotypes. Localities were obtained from museum records, and we used Google Earth to obtain geographic coordinates of the nearest forested area in the species range. Localities were listed as forests or towns with directional coordinates (e.g., 40 km W of Tamatave). We used R packages ‘raster’ and ‘sp’ to extract all localities’ climate data in the WorldClim database (Hijmans & Etten, 2012; Pebesma & Bavand, n.d.). We ran a correlation matrix for all 19 bioclimatic variables and removed any highly correlated variables (r > .80). In the end, we used five bioclimatic variables in our comparative analysis as predictor variables (SI 5).

### DNA extractions for opsin genotypes

We categorized diurnal Indriidae color vision status (e.g., polymorphic color vision, dichromatic – red-green colorblind) based on X-linked opsin genotypes (Jacobs et al., 2017, 2019). Our analysis included diurnal Indriidae represented by a subset of species/populations for which pelage data were also available (*N*_species_ = 7; *N*_populations_ = 10; SI 6). Opsin genotypes for populations of diademed sifaka (*Propithecus diadema*), Milne-Edwards’s sifaka (*Propithecus edwardsi*), golden crowned sifaka (*Propithecus tattersalli*), Verreaux’s sifaka (*Propithecus verreauxi*), and *Indri indri* were previously published (Jacobs et al., 2017). Data for the silky sifaka (*Propithecus candidus*) and Perrier’s sifaka (*Propithecus perrieri*) were based on opsin genotypes obtained from five individuals from Anjanaharibe-Sud (N_X chromosomes_ = 8) and Analamera (N_X chromosomes_ = 8) populations, respectively, following (Jacobs et al., 2019). Briefly, we extracted genomic DNA from blood/tissue samples using phenol-chloroform-isoamyl extraction protocols (Sambrook et al., 1989). We used quantitative PCR (Mic qPCR Cycler, Bio Molecular Systems; Rotor-Gene Q, Qiagen) to amplify exons 3 and 5 of the X-linked opsin gene. Amplification was followed by high-resolution melt analysis (HRMA) (*following* (Jacobs et al., 2016)), and we determined genotypes for each exon based on the observed melt curve shape and temperature compared to positive controls (N_Replicates_ = 2 per exon). To further confirm genotype calls, we Sanger sequenced qPCR amplicons from at least two individuals representing each observed melt curve. All sequences matched scored genotypes based on melt curves.

We estimated the peak spectral sensitivity of observed opsin alleles in each population (*following* (Jacobs et al., 2017)) and determined the difference between the two alleles with maximum and minimum peak spectral sensitivities. These values, along with a total number of observed alleles, were used in downstream analyses as proxies for enhanced trichromatic color vision. Trichromatic color vision is most optimal with more widely separated photopigments (e.g., (Osorio et al., 2004)). All else being equal, a greater number of opsin alleles results in an increased proportion of heterozygous (trichromatic) females in a population.

### Analyses

#### Climate and body size effects on pelage brightness and hue

We conducted four principal component analyses (PCAs): two across Indriidae and two within *Propithecus*. One was for all the brightness variables across body regions and a second for all the hue variables across body regions for Indriidae, to test specific hypotheses related to each and to reduce the dimensionality of the data. We then ran two additional albeit similar PCAs within only the *Propithecus* genus—given that the *Propithecus* genus is more widely geographically distributed than either *Avahi* or *Indri*. We removed any variables with strongly correlated vectors in the PCAs and then re-ran them to obtain the final PC values. We only used the resulting PC values with eigenvalues over one as our dependent variables.

To test the effects of climate on pelage color while accounting for the evolutionary relatedness of taxa, we used phylogenetic generalized linear mixed models (PGLMM) using Markov chain Monte Carlo algorithms (MCMC) via the ‘MCMCglmm’ package in R (Hadfield, 2010). Our Indriidae phylogeny was based on a previously published lemur phylogeny that contains a complete picture of Indriidae evolution (Herrera & Dávalos, 2016). We assigned the phylogeny to the ’pedigree’ argument in MCMCglmm, and the resulting matrix was used to weigh the error structure of the model. Similar methodologies have been used to test the effects of climate variables on bumblebee body size (Ramírez-Delgado et al., 2016) and frog vocal signals (McLean et al., 2013). Each model included one color PC value as the dependent variable and the bioclimatic variables (SI 5) and body size as independent variables, with species and phylogeny considered random factors. We log-transformed PC values, bioclimatic variables, and body size to achieve normality. All models ran for 5,000,000 iterations after a burn-in of 500,000 iterations and a thinning interval of 500 using flat priors. We assured effective sampling sizes were adequate for all our models (i.e., ESS virtually equal to MCMC sample size) (Villemereuil & Nakagawa, 2014).

We assigned statistical significance when pMCMC ≤ 0.05 (which is equivalent to a p-value (Villemereuil & Nakagawa, 2014)) and when the confidence interval for a given variable did not include zero (Ramírez-Delgado et al., 2016). To estimate what fraction of the pelage variation was attributable to phylogenetic relatedness, we calculated *H^2^* from Lynch (Lynch, 1991) (equivalent to Pagels’s λ) which corresponds to the degree of variation in a given trait attributable to phylogenetic relatedness (Hansen & Orzack, 2005; Kamilar & Cooper, 2013). We ran the models across the whole Indriidae family and within the *Propithecus* genus. Using G*Power analysis (Faul et al., 2009), we estimated the PGLMMs would detect a small effect ((Indriidae, 0.25)-(*Propithecus*, 0.35)) with ≈80% power.

#### Climate and body size effects on pelage density

We conducted two principal component analyses (PCA) of hair and follicle density values: one across all Indriidae taxa and a second within the *Propithecus* genus. We removed any variables whose vectors strongly correlated in each of the PCAs. We used the resulting PC values with eigenvalues over one as dependent variables in our PGLMM models, following the methodology outlined above (see: climate and body size effects on pelage brightness and hue).

Using G*Power analysis (Faul et al., 2009), we estimated the Indriidae PGLMM would detect a small effect (0.35), and the *Propithecus* PGLMM would detect a moderate effect (0.50) with ≈80% power. We also conducted an Analysis of Variance (ANOVA) to test for differences between hair or follicle density across distinct body parts.

#### Polymorphic trichromacy and pelage hue variation

We ran one PCA on hue variables among a subset of individuals from which we had paired opsin genotypes and pelage measurements from the same population. We used any PCs with eigenvalues over one as the dependent variable in our linear models. To estimate the relationship between pelage hue and color vision, we conducted both (1) a simple linear regression and (2) PGLMM. Due to the small sample size (n = 10 localities/populations) (SI 6), we assumed that the "true" result likely fell between a model where lambda = 0 (simple linear model) and one where lambda = 1 (PGLMM). We followed the methods described previously, except we used ‘spectral sensitivity range’ (i.e., difference in opsin spectral sensitivity) and the ‘total number of opsin alleles’ as our independent variables. Despite the low sample size, using G*Power analysis (Faul et al., 2009), we estimated the simple linear model would detect a moderate effect (0.65) with ≈80% power.

## Results

### Climate and body size effects on pelage brightness and hue

#### Across Indriidae

Across Indriidae, the first PC for the brightness PCA explained ≈61% of the variation (SI 7). The highest loadings on PC1 were hindlimb, tail, and cheek brightness (SI 8). We found a statistically significant and positive association between hair brightness and precipitation seasonality (Table 1). Thus, darker pelage across Indriidae is associated with forests of lower precipitation seasonality (i.e., higher rainfall, more humid) (Figure 2A). Brightness across Indriidae exhibited a moderate phylogenetic signal (0.68) (Table 1). On the hue PCA, the first three PCs explained ≈68% of the variation (SI 9, SI 10). The highest loadings were forelimb, hindlimb and ventral torso hue (PC1), cheek and ventral torso (PC2), as well as crown hue (PC3) (SI 10). We found a statistically significant relationship between PC1 and PC2 with precipitation seasonality (Table 2). Indriidae with redder pelage hues live in regions that experience less seasonality (i.e., higher rainfall) (Figure 3, SI 24). Hue values across Indriidae exhibited low to moderate phylogenetic signal (0.25-0.48) (Table 2).

**Figure 2.**
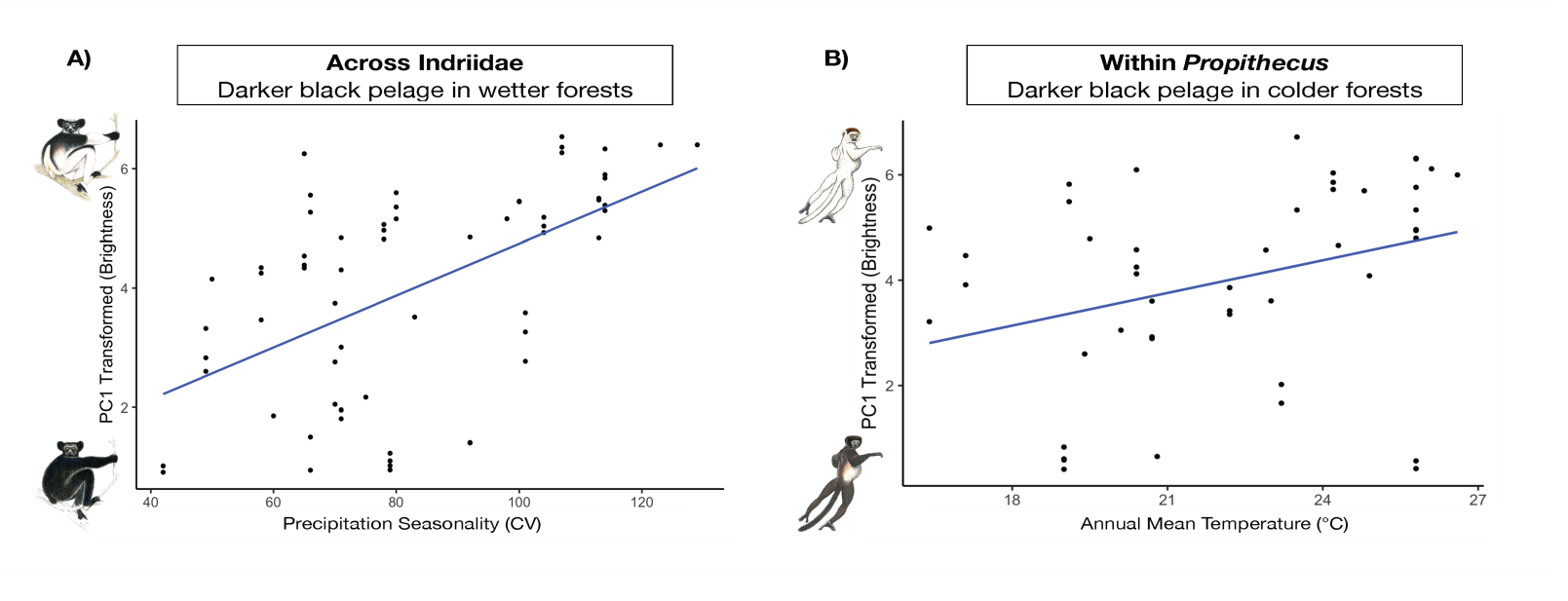
Plot of brightness across (A) the entire Indriidae family, in association with precipitation Seasonality (CV) (showing adherence to Gloger’s Rule), and (B) only within the *Propithecus* genus, in association with Annual Mean Temperature (°C) (showing adherence to Bogert’s Rule).

**Figure 3.**
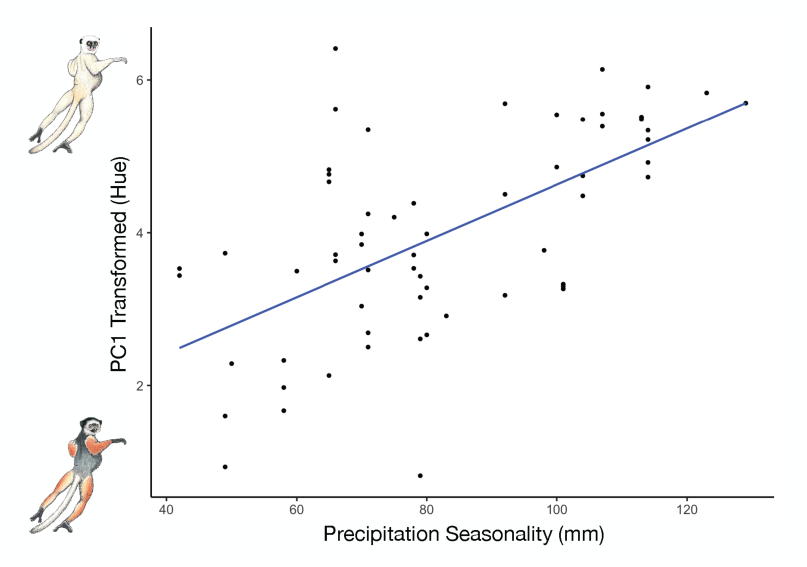
Plot of pelage hue trends across (A) the entire Indriidae family, in association with precipitation seasonality (mm) (showing adherence to Gloger’s Rule). Results are similar within *Propithecus*.

**Table 1.**
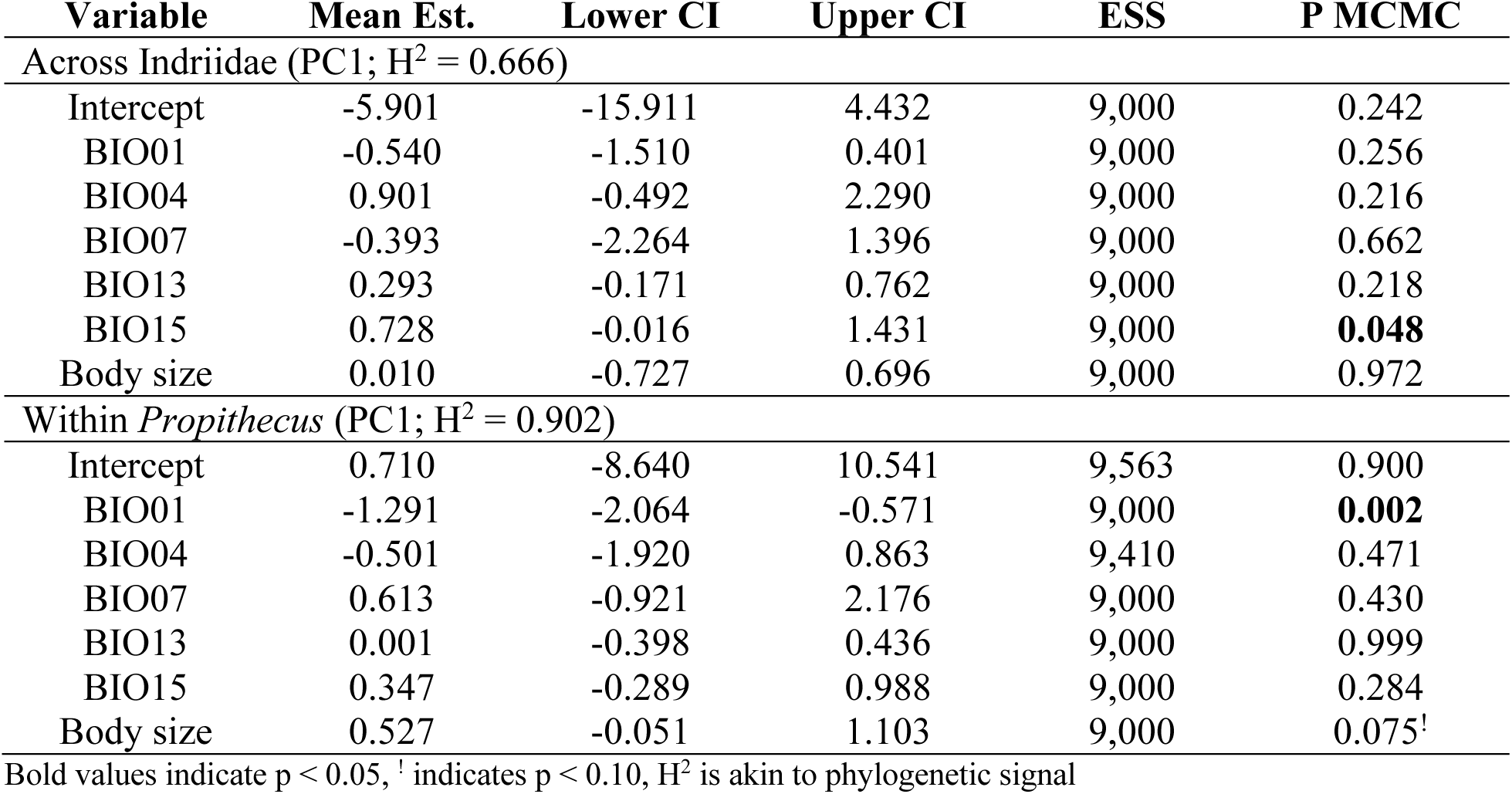
Results of PGLMMs predicting hair brightness (RGB averages) across all Indriidae taxa and only within *Propithecus*

**Table 2.**
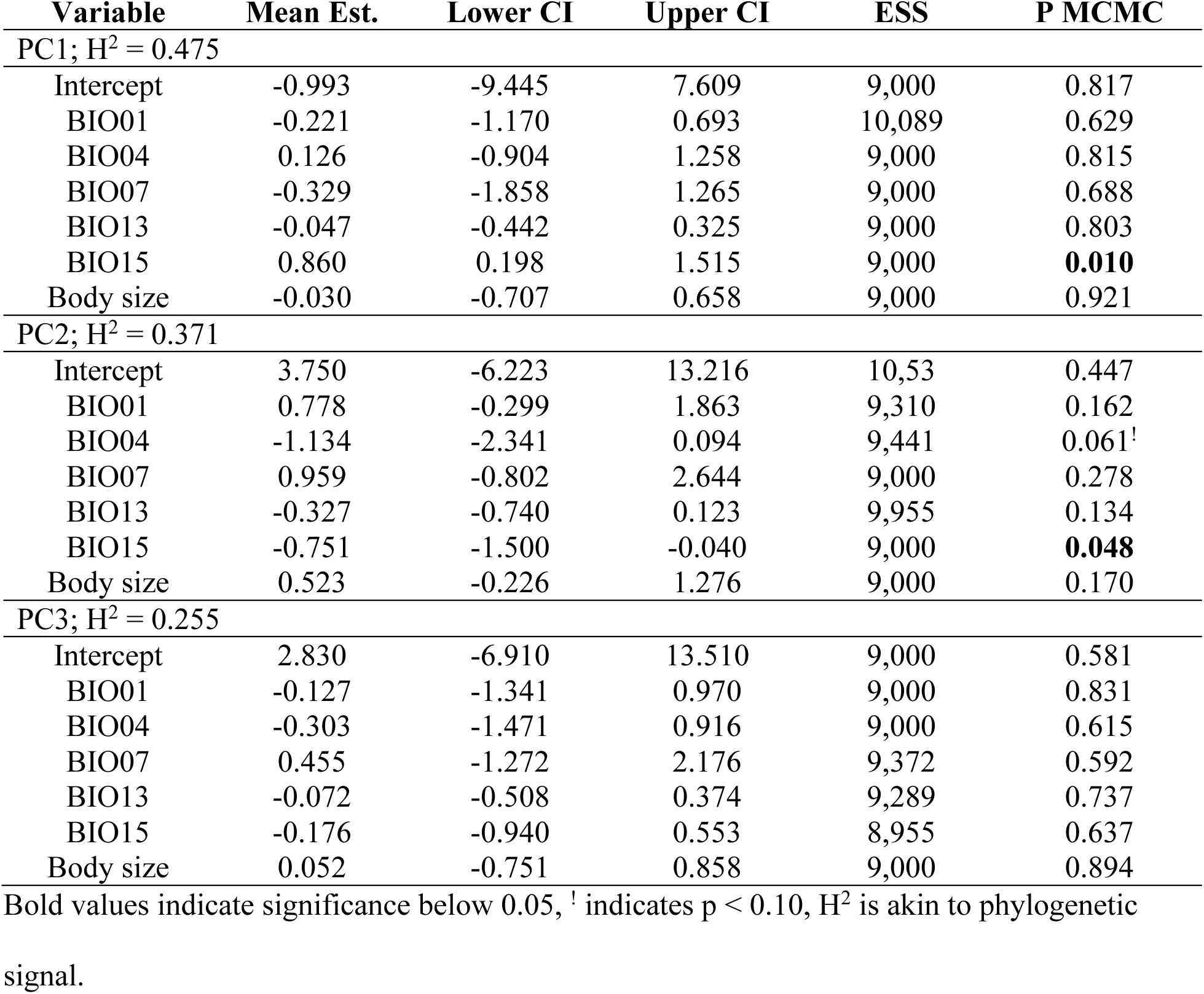
Results of PGLMMs predicting hair hue (RG ratio values) across all Indriidae taxa

#### Within *Propithecus*

For the brightness PCA, the first PC explained ≈70% of the variation within *Propithecus* (SI 11). The highest loadings on PC1 were hindlimb, dorsal torso, and forelimb brightness (SI 12). We found a statistically significant and positive association between hair brightness and mean annual temperature (Table 1). Thus, darker pelage within *Propithecus* is associated with colder forests (Figure 2B). Brightness across *Propithecus* exhibited a high phylogenetic signal (0.90) (Table 1). For the hue PCA, the first three PCs explained ≈80% of the variation within *Propithecus* (SI 13, SI 14). The highest loadings were forelimb, dorsal torso, and hindlimb hue (PC1), cheek, ventral torso, and hindlimb hue (PC2), and crown hue (PC3) (SI 14). We found a statistically significant relationship between PC2 and mean annual temperature, precipitation seasonality, precipitation of the wettest month, and body size (Table 3). Thus, redder hues within *Propithecus* are associated with lower precipitation seasonality (and higher rainfall). We found no statistically significant relationships between either PC1 or PC3 with the tested variables.

**Table 3.** Results of PGLMMs predicting hair hue (RG ratio values) within the *Propithecus* genus

| Variable | Mean Est. | Lower CI | Upper CI | ESS | P MCMC |
| --- | --- | --- | --- | --- | --- |
| PC1; H<sup>2</sup> = 0.632 |  |  |  |  |  |
| Intercept | 7.664 | -5.813 | 21.243 | 8,635 | 0.268 |
| BIO01 | -0.341 | -1.656 | 0.811 | 9,000 | 0.584 |
| BIO04 | -0.567 | -2.530 | 1.366 | 8,581 | 0.565 |
| BIO07 | 0.751 | -1.599 | 3.290 | 8,631 | 0.548 |
| BIO13 | 0.004 | -0.612 | 0.607 | 8,699 | 0.984 |
| BIO15 | -0.969 | -2.021 | 0.057 | 8,524 | 0.069<sup>†</sup> |
| Body size | -0.083 | -1.063 | 0.831 | 9,375 | 0.857 |
| PC2; H<sup>2</sup> = 0.513 |  |  |  |  |  |
| Intercept | 1.773 | -8.206 | 11.490 | 8,358 | 0.728 |
| BIO01 | 1.434 | 0.448 | 2.381 | 9,322 | **0.006** |
| BIO04 | -1.220 | -2.587 | 0.153 | 8,042 | 0.076<sup>†</sup> |
| BIO07 | 1.154 | -0.598 | 2.923 | 9,000 | 0.187 |
| BIO13 | -0.450 | -0.907 | -0.004 | 9,000 | **0.045** |
| BIO15 | -1.089 | -1.862 | -0.314 | 9,000 | **0.006** |
| Body size | 0.806 | 0.047 | 1.531 | 8,829 | **0.037** |
| PC3; H<sup>2</sup> = 0.227 |  |  |  |  |  |
| Intercept | 3.296 | -0.121 | 1.789 | 9,395 | 0.639 |
| BIO01 | -0.429 | -2.065 | 1.199 | 9,000 | 0.596 |
| BIO04 | -0.155 | -1.744 | 2.085 | 9,492 | 0.903 |
| BIO07 | -0.753 | -3.283 | 1.704 | 9,765 | 0.545 |
| BIO13 | -0.008 | -0.643 | -0.637 | 9,000 | 0.975 |
| BIO15 | 0.001 | -1.241 | 1.213 | 9,325 | 1.000 |
| Body size | -0.148 | -1.166 | 1.471 | 9,000 | 0.812 |

Bold values indicate significance below 0.05, <sup>†</sup> indicates p < 0.10, H<sup>2</sup> is akin to phylogenetic signal.

There was low to moderate phylogenetic signal for hair hue within *Propithecus* (0.22-0.63) (Table 3).

### Climate and body size effects on pelage density

#### Across Indriidae

Across Indriidae, the first three PCs explained ≈70% of the variation (SI 15, SI 16). The highest loadings were forelimb, hindlimb, and crown follicle density (PC1), ventral torso hair density and crown and hindlimb follicle density (PC2), and dorsal torso hair density (PC3) (SI 16). We found a statistically significant relationship between PC1 and precipitation seasonality as well as between PC3 and body size (Table 4). Specifically, the results suggest that in forests with high precipitation seasonality (dryer, open forests exposed to the sun) individuals exhibit an increase of follicle density on the limbs and crown (Figure 4). Also, smaller taxa have the highest hair density; and, larger individuals exhibit lower hair density on average (Figure 5). Pelage density across Indriidae exhibited low phylogenetic signal (0.21-0.33) (Table 4). We also found a statistically significant difference between hair density across body regions sampled (Table 5). Specifically, we found the crown and forelimbs exhibit the greatest hair density, and the ventral torso exhibits the least hair density (Figure 6A). However, we found no significant differences in follicle density of regions sampled (Table 5, Figure 6B).

**Figure 4.**
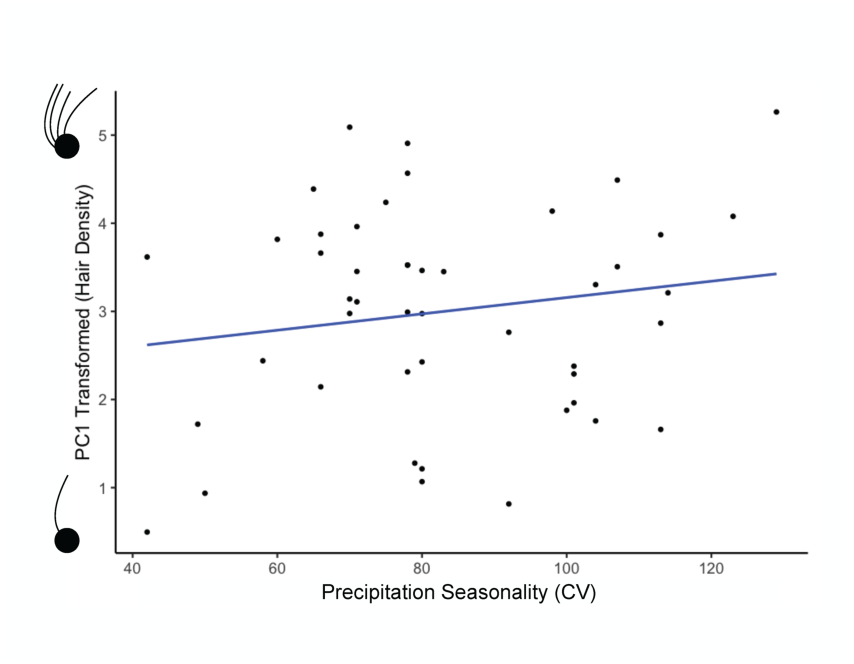
Plot of hair density trends. The relationship between PC1 and precipitation seasonality (CV) is significantly associated with hair density on the limbs and crown across the Indriidae family. Lower PC1 values are associated with lower hair density on the limbs and crown.

**Figure 5.**
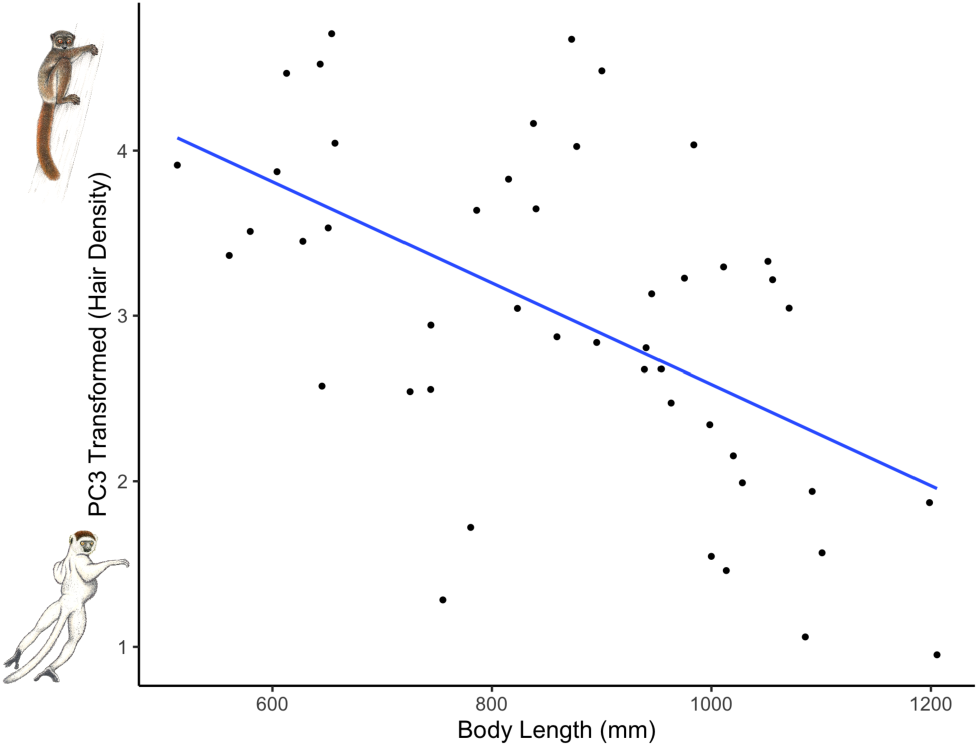
Plot of hair density trends. The relationship between PC3 and body length (capturing length, in mm) is significantly associated with hair density on the dorsal torso across Indriidae. Lower PC3 values are associated with lower hair density across the dorsal torso.

**Figure 6.**
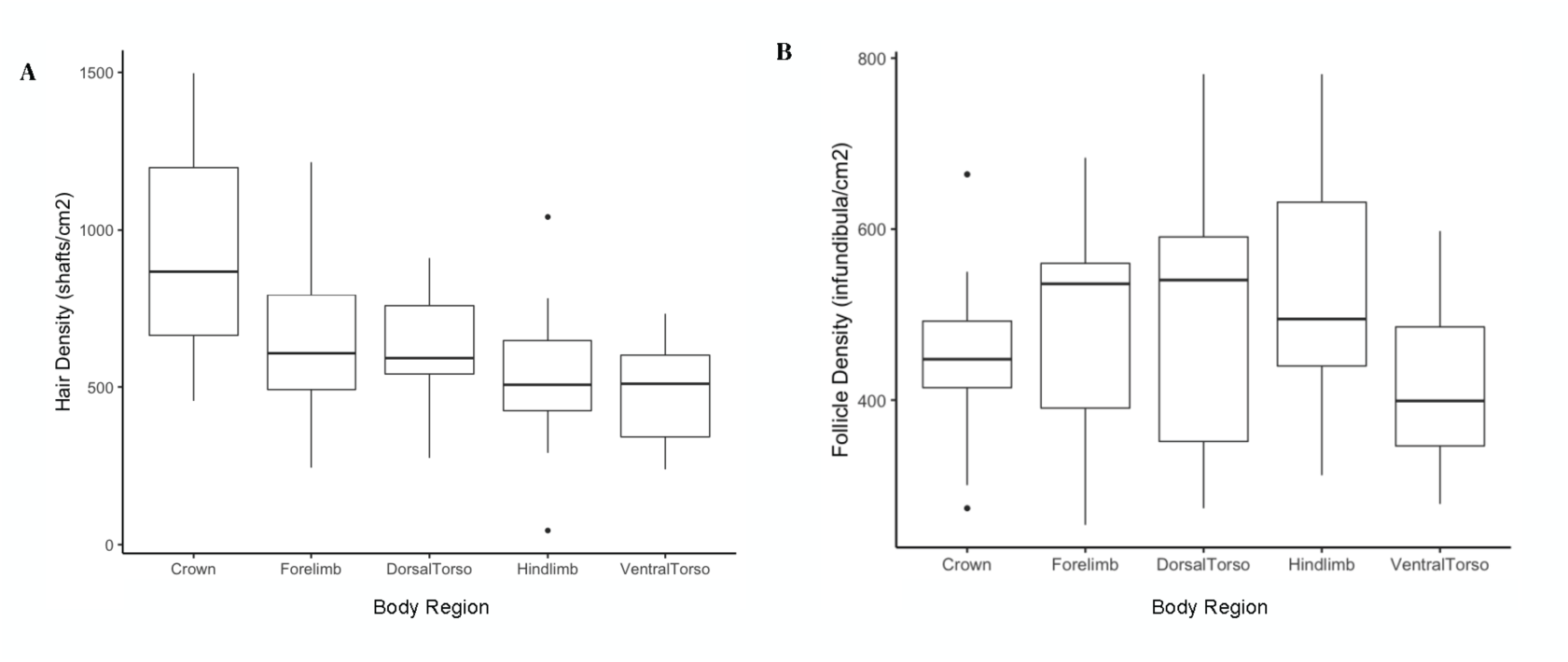
Boxplots illustrating variation in (A) hair density and (B) follicle density across body regions, for Indriidae. Black lines represent the average, and black dots represent outliers.

**Table 4.** Results of PGLMMs predicting hair and follicle density across all Indriidae taxa

| Variable | Mean Est. | Lower CI | Upper CI | ESS | P MCMC |
| --- | --- | --- | --- | --- | --- |
| PC1; H<sup>2</sup> = 0.222 |  |  |  |  |  |
| Intercept | -6.478 | -22.987 | 18.456 | 9,456 | 0.401 |
| BIO01 | -0.036 | -1.368 | 1.366 | 9,000 | 0.961 |
| BIO04 | 0.770 | -0.673 | 2.228 | 9,000 | 0.278 |
| BIO07 | -1.093 | -3.262 | 1.047 | 9,434 | 0.314 |
| BIO13 | 0.523 | -0.085 | 1.190 | 9,000 | 0.101 |
| BIO15 | 1.232 | 0.185 | 2.255 | 9,290 | **0.020** |
| Body size | -0.275 | -1.326 | 0.770 | 9,339 | 0.597 |
| PC2; H<sup>2</sup> = 0.217 |  |  |  |  |  |
| Intercept | 0.256 | -15.042 | 16.829 | 9,000 | 0.981 |
| BIO01 | 0.175 | -1.238 | 1.587 | 9,000 | 0.812 |
| BIO04 | 0.150 | -1.334 | 1.643 | 9,000 | 0.835 |
| BIO07 | -0.990 | -3.299 | 1.243 | 9,000 | 0.375 |
| BIO13 | 0.110 | -0.523 | 0.806 | 9,000 | 0.736 |
| BIO15 | 0.518 | -0.550 | 1.585 | 9,765 | 0.331 |
| Body size | -0.161 | -1.266 | 0.922 | 8,309 | 0.760 |
| PC3; H<sup>2</sup> = 0.330 |  |  |  |  |  |
| Intercept | 10.259 | -0.399 | 20.576 | 9,000 | 0.057<sup>†</sup> |
| BIO01 | -0.259 | -1.175 | 0.700 | 9,000 | 0.602 |
| BIO04 | 0.071 | -0.953 | 1.056 | 9,307 | 0.876 |
| BIO07 | -0.129 | -1.643 | 1.413 | 9,000 | 0.853 |
| BIO13 | -0.337 | -0.772 | 0.081 | 9,000 | 0.123 |
| BIO15 | 0.438 | -0.276 | 1.164 | 9,908 | 0.228 |
| Body size | -1.265 | -2.012 | -0.514 | 9,000 | **0.002** |

Bold values indicate significance below 0.05, <sup>†</sup> indicates p < 0.10, H<sup>2</sup> is akin to phylogenetic signal.

**Table 5.** Summary of ANOVA examining hair and follicle density variation across distinct body regions for Indriidae

|  | DF | Sum Sq. | Mean Sq. | F value | P value |
| --- | --- | --- | --- | --- | --- |
| *Hair density* |  |  |  |  |  |
| Body Region | 4 | 1458536 | 364634 | 5.808 | **< 0.0005** |
| Residuals | 60 | 3766879 | 62781 |  |  |
| *Follicle density* |  |  |  |  |  |
| Body Region | 4 | 108592 | 27148 | 1.524 | 0.207 |
| Residuals | 60 | 1068659 | 17811 |  |  |

#### Within *Propithecus*

Within *Propithecus*, the first three PCs explained ≈70% of the variation (SI 17, SI 18). The highest loadings were forelimb follicle density, crown hair density, and hindlimb follicle density (PC1), crown and hindlimb follicle density as well as hindlimb hair density (PC2), and ventral torso hair density (PC3) (SI 18). We found a statistically significant relationship between PC1 and precipitation seasonality, similar to Indriidae (Table 6). However, unlike the Indriidae results, none of the PCs were significant with body size. Though, we detected significant differences between hair density and distinct body regions (SI 19), and no differences were detected among follicle density within *Propithecus* (SI 19, SI 20).

**Table 6.**
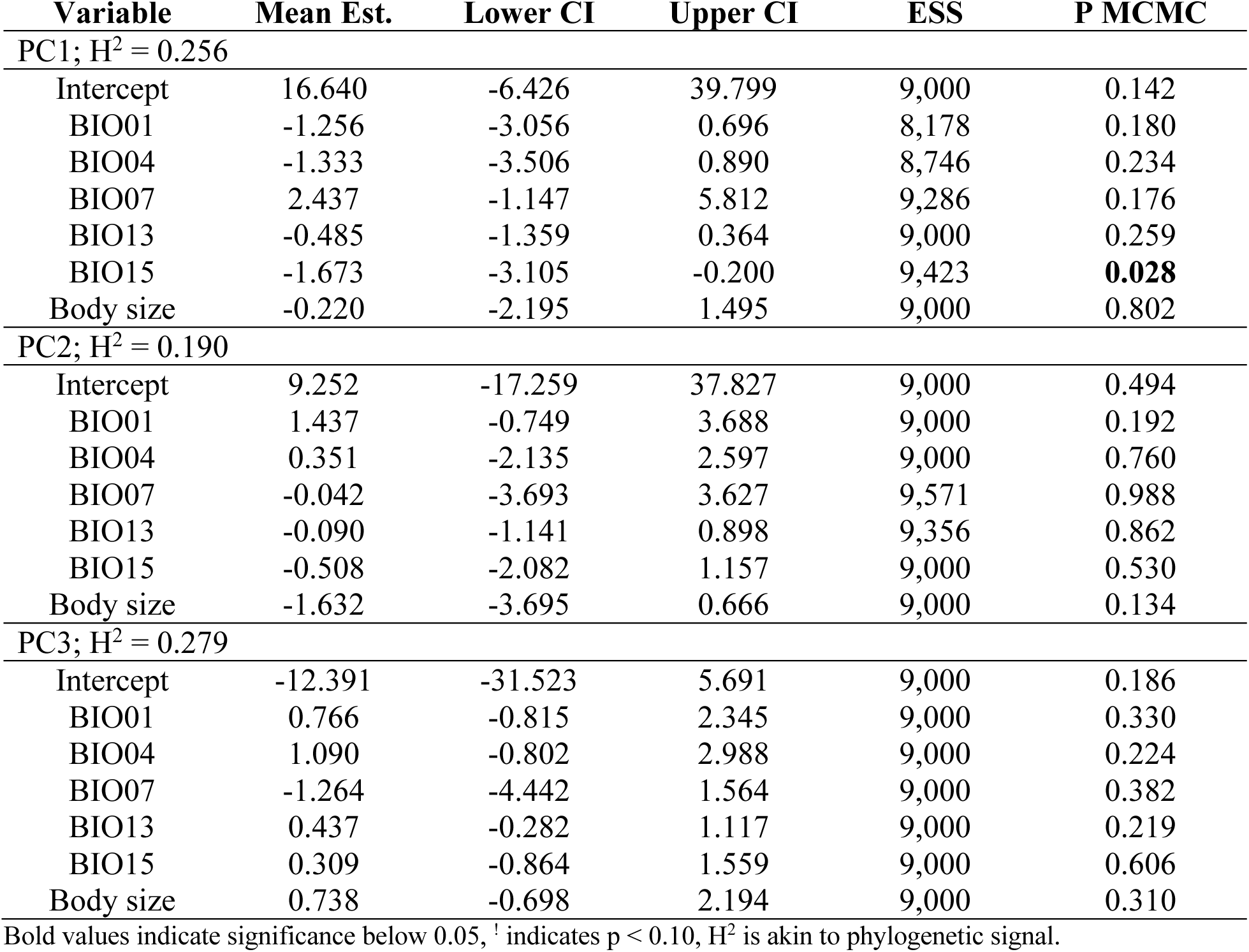
Results of PGLMMs predicting hair and follicle density within the *Propithecus* genus

### Polymorphic trichromacy and pelage hue variation

For the hue PCA using only a subset of the data from *Indri* and *Propithecus*, the first two PCs explained ≈66% of the variation (SI 21). The highest loadings to PC1 were forelimb, dorsal torso, and hindlimb hue, and the highest loadings to PC2 were cheek, ventral torso, and hindlimb hue (SI 22). From the simple linear model (lambda = 0), we found a statistically significant relationship between PC1 with the difference in opsin spectral sensitivity (p = 0.03; R^2^ = .51), but not with the overall number of alleles in the population (Table 7). Specifically, we found a significant negative linear relationship between spectral sensitivity and hue (Figure 7). Despite the smaller sample size of the PGLMM (lambda = 1), we detected a similar association that approaches statistical significance (pMCMC = 0.09) and only moderate phylogenetic signal (0.41) (SI 23).

**Figure 7.**
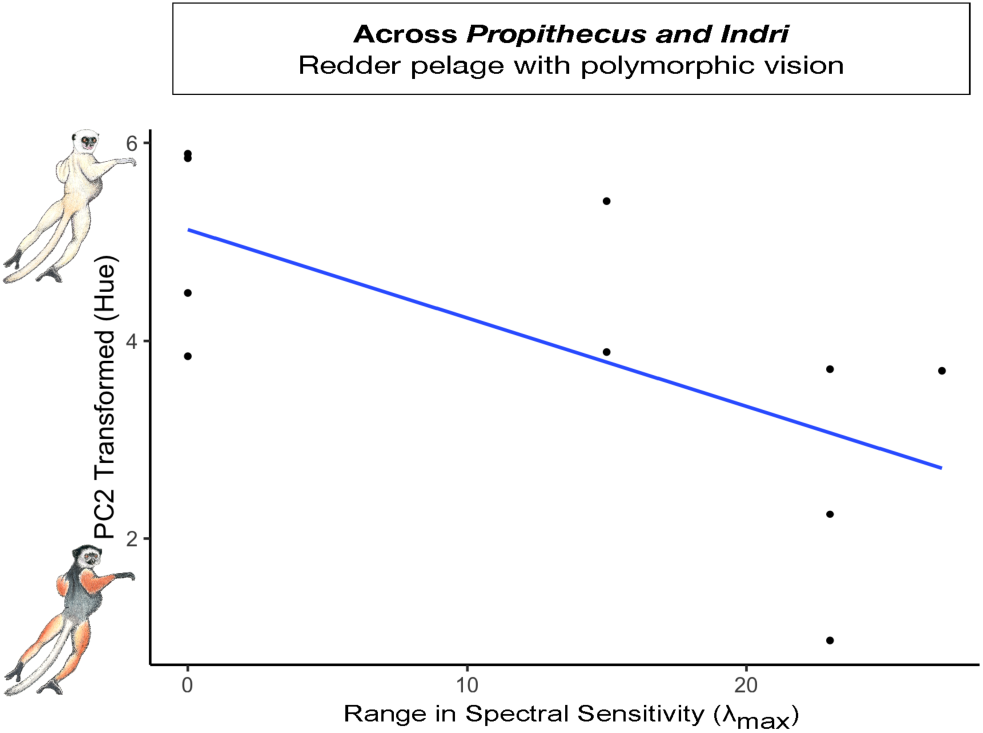
Plot of pelage hue trends across diurnal *Propithecus* and *Indri* genera, in association with the range (i.e., difference) in opsin peak spectral sensitivity.

**Table 7.** Summary of linear models examining pelage coloration against difference in opsin peak spectral sensitivity, using hue (PC1) scores of individuals across a subset of 10 sampling locations

|  | Estimate | Std. Error | T value | P value |
| --- | --- | --- | --- | --- |
| PC1 |  |  |  |  |
| [Intercept] | -1.529 | 1.611 | -0.949 | 0.374 |
| Difference in Spectral Sensitivity | -0.234 | 0.089 | -2.630 | **0.034** |
| Number of Opsin Alleles | 2.635 | 1.501 | 1.746 | 0.124 |

SE: 1.05 on 7 degrees of freedom; $R^2 = 0.516$ , $F = 5.801$ on 2 and 7 degrees of freedom; $p = 0.033$ . Bold values indicate significance below 0.05.

## Discussion

Our overarching goals were to examine 1) if Indriidae pelage traits varied across distinct phylogenetic scales and 2) if multiple non-exclusive selective pressures would impact the same trait. We found support for both; distinct aspects of pelage color and density in Indriidae and sifakas (genus *Propithecus*) vary with climate, body size, and/or color vision.

### Climate and body size effects on pelage brightness and hue

Our first sub-aim sought to uncover how climate and body size influenced pelage brightness and hue across two distinct scales (across a family [Indriidae] and within a genus [*Propithecus*]). Our results show distinct trends across scales regarding dark pelage (i.e., brightness). Specifically, across Indriidae, greater brightness is associated with greater precipitation seasonality (Table 1). Greater hue, similarly, is associated with less precipitation seasonality across Indriidae and *Propithecus* (SI 24)) (Table 2). Thus, more pigmented coats (i.e., individuals with blacker and redder hair) occur in regions of lower precipitation seasonality (Figure 2A, Figure 3). Generally, in Madagascar, low precipitation seasonality is strongly and negatively associated with high precipitation forests of the East (SI 24). The result for brightness follows Gloger’s rule, predicting an increase in eumelanin (i.e., black pigments) (Delhey, 2017). Yet, within *Propithecus*, results indicate a relationship between brightness and mean annual temperature (Table 1). Thus, dark-haired sifakas are generally associated with colder regions, which serves as evidence of Bogert’s rule (or the thermal melanism hypothesis) (Figure 2B) (Bogert, 1949).

Recent work highlights that discovering evidence for Bogert’s or Gloger’s rule may merely reflect differences in geographic space, scale, and climate gradients (Delhey et al., 2019). We show in this analysis that different effects can be detected across the primate coat, depending on the sampling scale. Additionally, the strength of selective pressures acting on pelage may also play a role in detecting either rule. For example, if the need to camouflage or avoid parasites is high, rainfall (which predicts Gloger’s rule) may strongly affect pelage color (da Silva et al., 2016). This is because higher rainfall may also be associated with higher loads of parasites (Shearer & Ezenwa, 2020) or may indicate low light forested environments that enhance crypsis for dark animals (Caro, 2005; Delhey, 2018). In contrast, if the need for thermoregulation is high, temperature (which predicts Bogert’s rule) may have a more substantial effect on pelage color than rainfall (Galván et al., 2018). It is hypothesized that black colors aid with thermoregulation in colder climates since they likely increase absorption of ultraviolet rays. While this has been documented widely in ectoderms and recently in birds (Bishop et al., 2016; Bogert, 1949; Clusella Trullas et al., 2007; Delhey et al., 2019; Xing et al., 2016), to date, our results serve as the first empirical evidence for Bogert’s rule in mammals. Notably, the range of temperatures occupied by *Propithecus* species (≈10-27°C) corresponds exceptionally well to the range at which Bogert’s rule is expected to influence plumage color (≈10-25°C) in passerines (Delhey et al., 2019). At this temperature range there is much nuance in the climatic space between temperature and precipitation, which is in turn is hypothesized to influence the rate of diversification and maintenance of species barriers (Delhey et al., 2019; Martin et al., 2010). Insights from other primate species may further elucidate the importance of Bogert’s and Gloger’s rules to broader primate evolution.

Both body size and phylogeny seem to have minimal impact on pelage color. First, body size only influenced hair hue across the *Propithecus* clade and otherwise did not affect our coloration models across either scale (Table 2, Table 3). At first glance, smaller individuals seem to exhibit redder hues; however, this trend is driven mostly by *Propithecus* species on the western side of the island. This may indicate subtler variation between species that our analysis does not capture. However, these results may also reflect broader climatic trends, variations in color vision (addressed below), or is perhaps the product of our body size sampling methodology. Second, most of the coloration traits we sampled exhibited low to moderate phylogenetic signals. Our brightness models, on the other hand, had a high phylogenetic signal (≈0.80). Thus, brightness may implicate genetic drift (Losos, 2008; Revell et al., 2008). However, it is well established that primate coat color exhibits little to no phylogenetic signal and is a poor marker of phylogeny (Bell et al., 2021). An alternative explanation for such a high signal is that it is an “artificial” result from taxonomic inflation that merely reflects patterns of geographic variation (Kamilar & Muldoon, 2010).

### Climate and body size effects on pelage density

Our second sub-aim sought to uncover how climate and body size influenced pelage hair and follicle density across two distinct scales (across a family [Indriidae] and within a genus [*Propithecus*]). We predicted that body size would affect density because these variables are associated across primates. Body size may also have been one of the factors influencing the reduction of the hairy coat in hominins (Sandel, 2013; Schwartz & Rosenblum, 1981). We found support for this prediction across Indriidae (Table 4). Specifically, higher hair density is associated with smaller body sizes (*Avahi*), and lower hair density is found in larger animals across Indriidae (Figure 4). One explanation for this result may be that interfollicular distance increases as surface area increases, thus, decreasing hair density in larger animals (Sandel, 2013). In addition, this relationship may provide thermoregulatory benefits since larger animals may struggle more to dissipate heat (Schwartz & Rosenblum, 1981). Our models show that climate also influences pelage variation across Indriidae. Limb and crown follicle density increases as precipitation seasonality increases (i.e., shifting to dry open habitats) (Table 4; Figure 4). Interestingly, the crown and forelimbs are also the regions that exhibit the highest degree of hair density (Figure 6A)—but not follicle density (Figure 6B). Thus, there are likely two mechanisms to increase density: (1) increase hair follicle density or (2) increase the number of hairs emerging from each follicle. Since follicle density is determined in early embryogenesis, follicle density may be more strongly genetically determined than hair density (which may be more plastic to rapid environmental changes, such as seasonality) (Tapanes et al., 2021). Alternatively, follicle density may be easier to measure without error and could be a more reliable estimate for ‘hairiness.’

Nevertheless, the shift towards drier habitats exposed to more sun (such as Western spiny thickets) leads to an increase in follicle density in the limbs and crown (Figure 4), where hair density is already the highest of all regions (Figure 6A). Since all Indriidae are vertical clingers and leapers, the crown and upper limbs are the most directly impacted by the sun’s rays—as they are known to hold up their arms while jumping bipedally. Thus, increased hair density in these regions may aid individuals in this clade with protection from ultraviolet rays (and/or protect the underlying skin from abrasion in spiny thickets). We consider this density pattern may help prevent photolysis of folate, an essential metabolite in the formation of the embryonic neural tube. Protection against folate leaching was likely a selection target in the evolution of high melanization within the skin of humans living in high UV areas (Jablonski & Chaplin, 2000). In humans, lower levels of folate (or folate deficiency) are also often significantly associated with hair loss (Almohanna et al., 2019; Ertug, 2018), and folate may play a significant role within the hair follicle (Durusoy et al., 2009). Since the skin of most Indriidae is already highly melanized, it is not surprising that populations exposed to hotter environments and more direct sun often have lighter coats (Figure 2B, Figure 3)—potentially to prevent overheating due to lack of sweat glands. We note that although *Avahi* is nocturnal relative to *Propithecus* and *Indri*, recent work explains their activity patterns may be more accurately described as “cathermality sensu *lato.*” Mainly this is because certain species are active across daylight hours, depending on the lunar phase (Campera et al., 2019). Therefore, implications of the potential function of the pelage pattern detected across the family is not solely relegated to *Propithecus* and *Indri*. However, we note that other aspects of hair morphology that we did not measure (length, curvature) may influence the strength of this selective pressure differently in distinct genera (Lasisi, 2021). Future studies should consider smaller phylogenetic scales, and other aspects of hair morphology (e.g., length) and climate (e.g., wind speed). Nonetheless, the pattern that emerges from this data may signal that variation in hair density (i.e., higher hair density in the scalp) across the human body may have occurred early on as an early mechanism to prevent folate leaching from key body regions exposed to direct UV rays. This body hair pattern potentially preceded the evolution of “naked” skin and sweat glands.

Our data is also rare evidence supporting the body cooling hypothesis—from the perspective of hair density in non-human primates. The body cooling hypothesis (Wheeler, 1985; 1992) argued that bipedal hominins evolved naked skin to allow for evaporative cooling. However, Wheeler (1985, 1992) also argued as part of this hypothesis that bipedal hominins likely retained hair on the crown and shoulders to protect those areas from UV exposure. Indriidae, presenting similar postural locomotion to humans, exhibit follicle density increases in their body regions most exposed to potential UV damage. Indriidae present a unique opportunity to comparatively study potential tradeoffs between folate levels, hair density, hair color, and thermoregulation across and within populations to clarify potential mechanisms driving this pattern of hair variation in upright primates.

Our results indicate that body size is also likely contributing to hair density variation (Figure 5). However, the smallest primates within the Indriidae clade are also nocturnal; larger-bodied Indriidae are diurnal. Thus, “body size” is likely confounded by activity cycle. Our results show that within *Propithecus* alone, there was no variation in hair density detected with body size—but there was with climate (Table 6). This may be due to a smaller sample size available for *Propithecus* density which we estimated would not pick up small effects (G* power analyses detailed in methods). Alternatively, at smaller phylogenetic scales, other aspects of hair morphology that we did not measure (e.g., overall shape, length, thickness) may directly relate to body size best at a smaller phylogenetic scale. For example, there are no differences in hair density between chimpanzees and humans, but there are significant differences in gland density, sweat production, and overall hair morphology (Kamberov et al., 2018). Hair and follicle density was likely one of a few aspects of hair evolution that were potentially malleable and allowed rapid adaptation to changing climatic environments for hominins. More research is needed across non-human and human primate hair to understand why and how humans retained hair in the places they did (e.g., scalp, pubic)—a critical part of the story in the evolution of our apparent “nakedness.”

### Polymorphic trichromacy and pelage hue variation

Our third sub-aim sought to understand the relationship between color vision and pelage hue in diurnal genera. We found support for our prediction that populations with enhanced capacity for trichromatic color vision would also contain individuals with redder pelage (*as hypothesized in* (Sumner & Mollon, 2003)). Many primates are unique among placental mammals in having trichromatic color vision, a feature that is thought to be advantageous for foraging on "red" foods (i.e., ripe fruit, young leaves), detecting predators, and/or tracking conspecifics (Kawamura et al., 2012; Surridge et al., 2003). Indeed, reddish skin coloration observed in many primate species is thought to play a salient role in conspecific signaling for primates (Moreira et al., 2019). However, previous support for the coevolution of red hair and trichromacy in primates is mixed (Fernandez & Morris, 2007; Kamilar et al., 2013). Our results indicate a significant positive relationship between hue (’redness’) and measures of enhanced trichromacy in Indriidae (Table 7). The similarity in results between both models (lambda=0 vs. lambda=1) indicates a likely significant relationship between hue and opsin genotype variation in diurnal Indriidae. Specifically, red hues are more likely to occur when there is a larger difference in spectral sensitivity between opsin alleles (Figure 7), suggesting increased discriminability of such hues. Results across Indriidae, which exhibits more subtle inter- and intra-species variation in trichromacy than has been previously considered, suggests that there may be a more complex and nuanced coevolutionary relationship between red hair coloration in primates and trichromatic color vision that deserves further attention (Table 7).

### Conclusions

Climate, body size, and color vision all impact hair variation across Indriidae, but the extent that each exerts its influence on the phenotype varies by sampled phylogenetic scale. These results have considerable implications for our understanding of primate (including human) hair evolution because it suggests that larger macro-scale trends detected across primates may not sufficiently explain variation within a genus, species, or population. Understanding wild non-human primate hair can also potentially help shed light on the evolution of hominin hair. Further work on primate hair should sample across smaller geographic or phylogenetic scales and from diverse non-human and human populations.

## Acknowledgments

We thank the American Museum of Natural History, the Smithsonian National Museum of Natural History, and Harvard University’s Museum of Comparative Zoology. Special thanks to museum staff that helped with *Propithecus* collections, including Ian Tattersall, Darrin Lunde, Mark Omura, Eleanor Hoeger, and Marisa Surovy. We also thank the Government of Madagascar (Ministry of the Environment and Sustainable Development), SADABE, Madagascar Biodiversity Partnership (MBP), Omaha Zoo, the University of Antananarivo, and Anthropobiologie et Développement for facilitation and partnership. We thank members of GW PGL and Utah PEGL labs for relevant discussions and feedback on ideas presented here. Illustrations copyright 2020 Stephen D. Nash/IUCN SSC Primate Specialist Group, used and modified with permission. The findings and conclusions in this article are those of the author(s) and do not necessarily represent the views of the U.S. Fish and Wildlife Service. The work was supported by the Leakey Foundation, the Wenner Gren Foundation, the National Science Foundation (BCS #1546730, BCS #1606360), the George Washington University, Northern Illinois University, National Geographic CRE, Eppley Fund for Research, Primate Conservation Inc., and IdeaWild. Field work to collect biological samples was supported by Conservation International, Margot Marsh Biodiversity Foundation, and The Ahmanson Foundation. The funders had no role in study design, data collection and analysis, decision to publish, or preparation of the manuscript.

