## Supplementary material for "Hair phenotype diversity across Indriidae lemurs": S1

**Supplementary Files**

**SI 1** Total sample used in this study for pigmentation MCMCglmm analysis. Numbers in parenthesis indicate when and where lower specimen counts were available for the hair/follicle density MCMCglmm analysis.

| Species | $N_{Localities}$ | $N_{Individuals}$ |
|---|---|---|
| *Avahi laniger* | 3 (2) | 4 (2) |
| *Avahi meridionalis* | 1 | 1 |
| *Avahi occidentalis* | 1 | 2 (1) |
| *Avahi ramanantsoavana* | 1 | 3 |
| *Indri indri* | 3 | 6 |
| *Propithecus candidus* | 1 | 2 (1) |
| *Propithecus coronatus* | 1 | 3 |
| *Propithecus deckenii* | 3 | 5 (4) |
| *Propithecus diadema* | 7 (6) | 12 (8) |
| *Propithecus edwardsi* | 2 | 5 |
| *Propithecus perrieri* | 1 | 2 (1) |
| *Propithecus tattersalli* | 1 | 1 |
| *Propithecus verreauxi* | 8 | 17 (11) |
| TOTAL | 33 | 63 |

Key: Hair Density, $N_{HairDensityLocalities}$ = 31 localities, $N_{HairDensityIndividual}$ = 47 individuals

**SI 2** Sample origins for hair color and density morphology measurements

| Location | N |
|---|---|
| *Museum:* Harvard's Museum of Comparative Zoology (MCZ) | 18 |
| *Museum:* American Museum of Natural History (AMNH) | 32 |
| *Museum:* Smithsonian National Museum of Natural History (USNM) | 9 |
| *Wild:* Tsinjoarivo forest, Madagascar | 4 |
| TOTAL | 63 |

**SI 3** Example of how sampling was conducted when a region was divided into three parts using the 'ruler' tool in Photoshop to measure (A) the whole dorsal surface from the base of the neck to the base of the tail (length: 1491), that was then divided into (B) upper, (C) middle, and (D) lower.

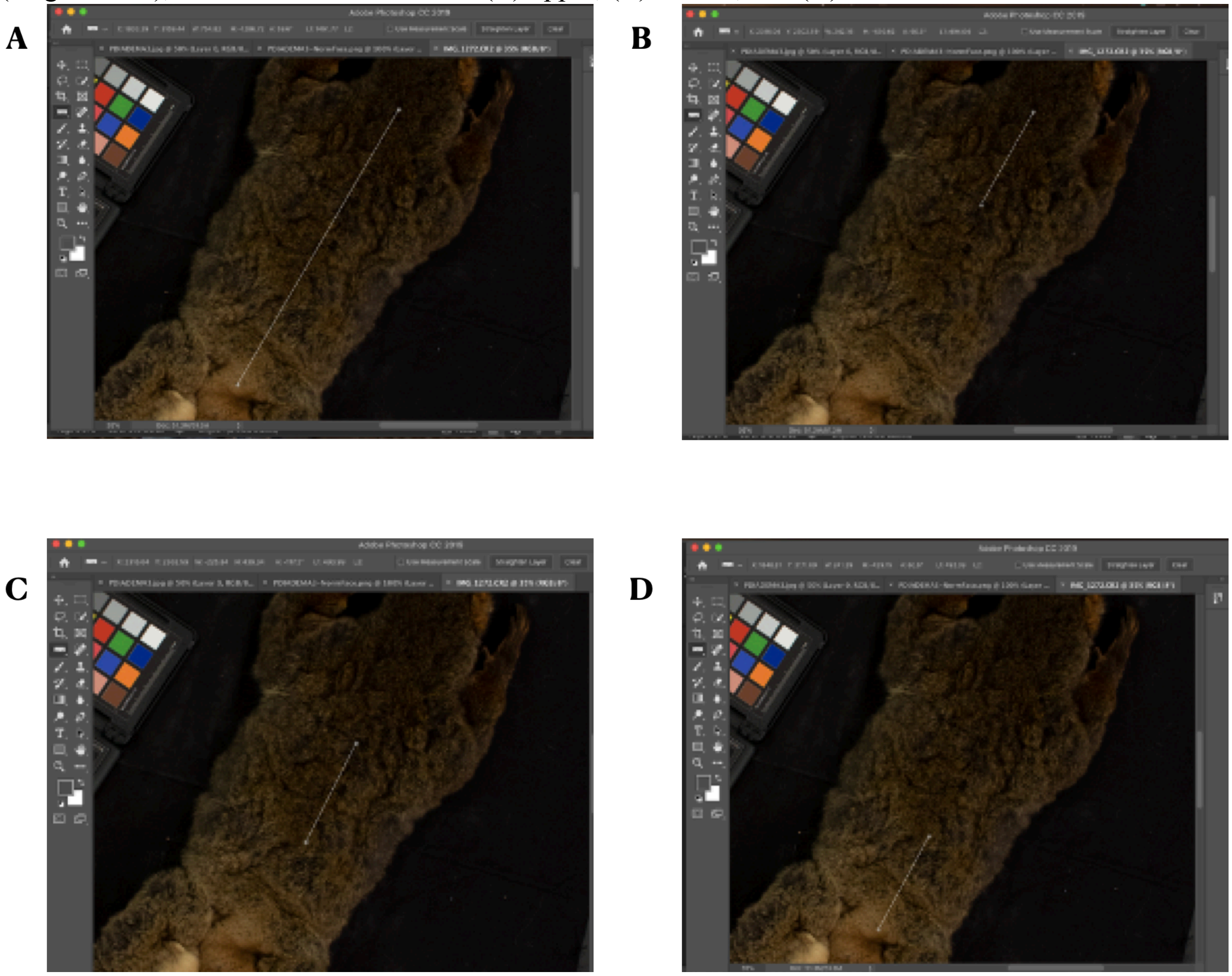

**SI 4** Schematic of the five body regions that were sampled for hair and follicle density

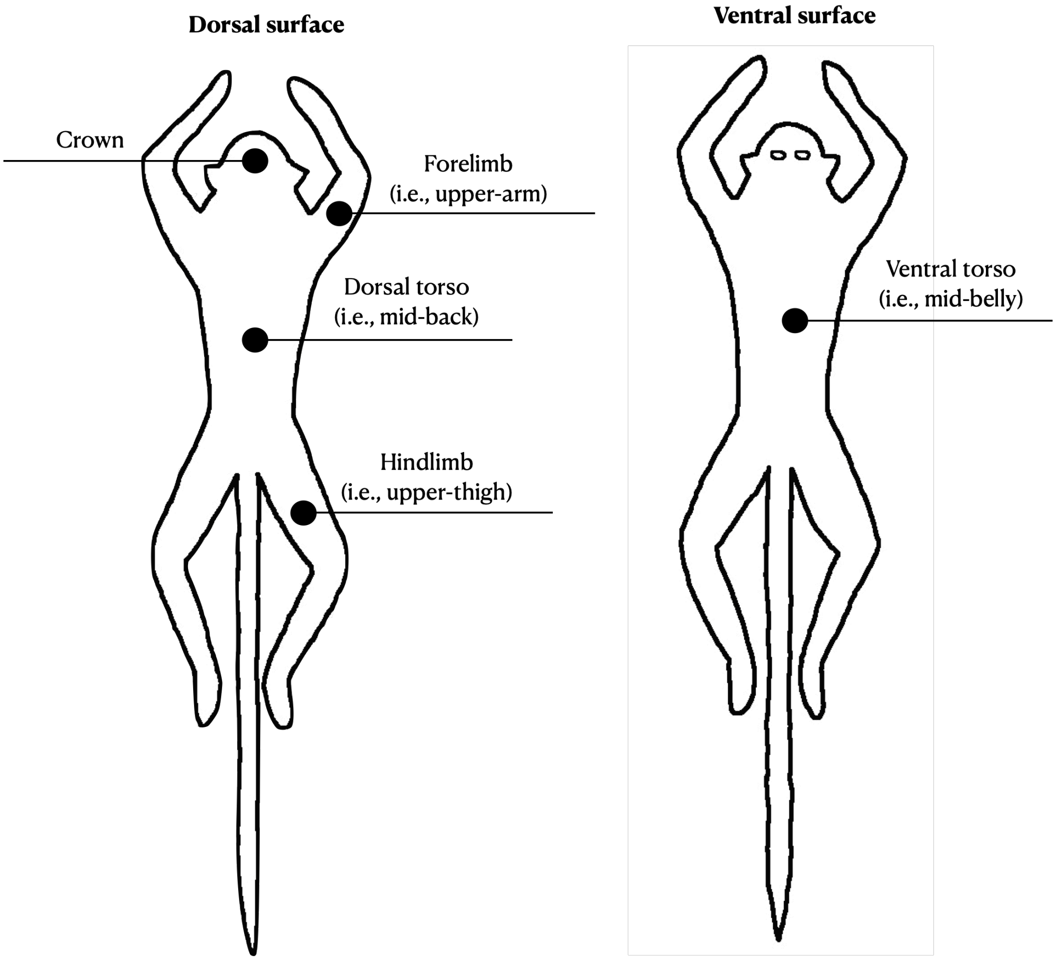

**SI 5** WorldClim bioclimatic variables used in the MCMCglmm analysis

| Layer Code | Bioclimatic Variable | Units |
|---|---|---|
| BIO 01 | Mean Annual Temperature | °C |
| BIO 04 | Temperature Seasonality (std. dev. x 100) | % |
| BIO 07 | Temperature Annual Range (BIO 05 – BIO 06) | °C |
| BIO 13 | Precipitation of Wettest Month | mm |
| BIO 15 | Precipitation Seasonality (Coefficient of Variation) | mm |

For more detailed descriptions, visit: http://worldclim.org/bioclim

**SI 6** Total sample statistics for samples used in opsin-pelage analysis

| Species | $N_{Localities}$ |
|---|---|
| *Indri indri* | 1 |
| *Propithecus candidus* | 1 |
| *Propithecus diadema* | 2 |
| *Propithecus edwardsi* | 1 |
| *Propithecus perrieri* | 1 |
| *Propithecus tattersalli* | 1 |
| *Propithecus verreauxi* | 3 |

Key: $N_{Localitites}$ = total number of locations from where we have pelage samples from.

**SI 7** PCA biplot of pelage brightness across genera for the Indriidae family

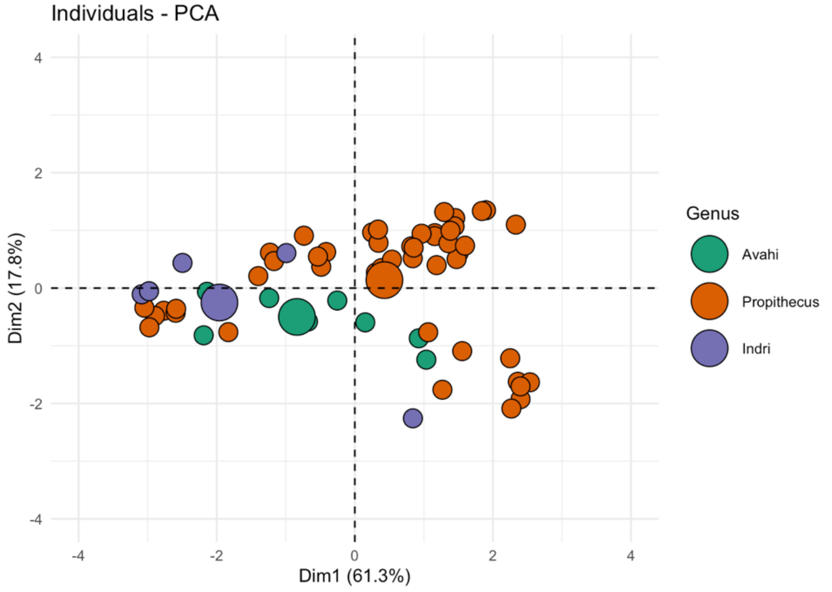

**SI 8** Loadings from the hair brightness PCA across Indriidae for each PC with an eigenvalue above one, from greatest to least variance explained.

| Principal Component | Variable | Variance Explained (%) |
|---|---|---|
| PC1 (eigenvalue = 1.96) | Hindlimb | 26.93 |
| PC1 | Tail | 22.75 |
| PC1 | Cheek | 22.68 |
| PC1 | Ventral torso | 19.66 |
| PC1 | Crown | 7.97 |

Contribution to overall variance: PC1 (61.26%). Total variance explained by PCs with eigenvalues > 1 is 61.26%.

**SI 9** PCA biplot of pelage hue across genera for the Indriidae family

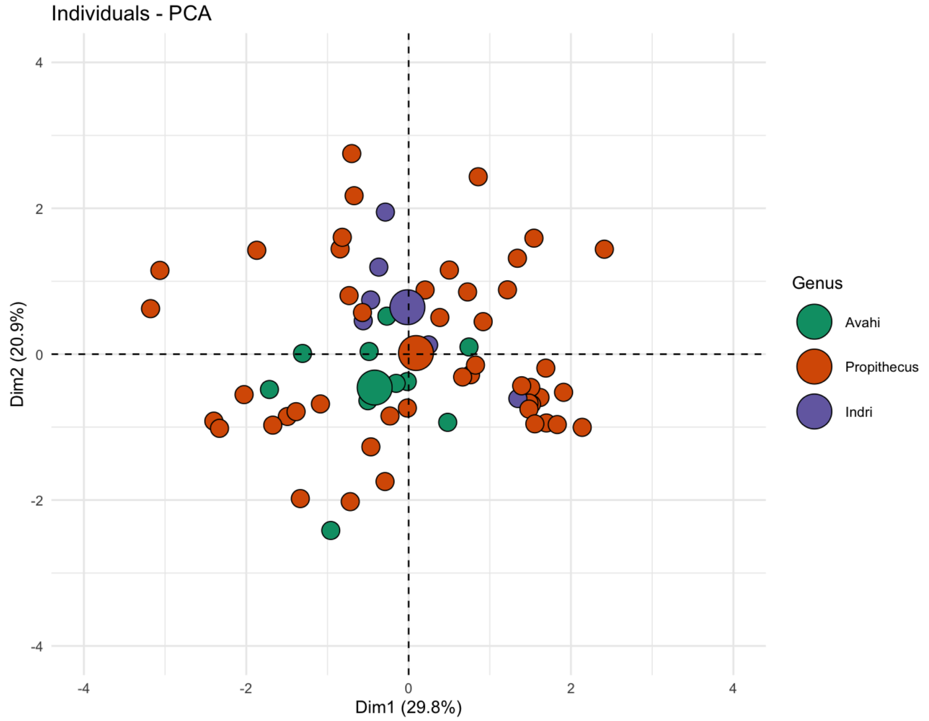

**SI 10** Loadings from the hair hue PCA across Indriidae for each PC with an eigenvalue above one, from greatest to least variance explained.

| **Principal Component** | **Variable** | **Variance Explained (%)** |
|---|---|---|
| PC1 (eigenvalue = 1.34) | Forelimb | 42.91 |
| PC1 | Hindlimb | 38.59 |
| PC1 | Ventral Torso | 11.67 |
| PC1 | Cheek | 5.03 |
| PC1 | Dorsal Torso | 1.27 |
| PC1 | Crown | 0.55 |
| PC2 (eigenvalue = 1.12) | Cheek | 40.56 |
| PC2 | Ventral torso | 35.47 |
| PC2 | Hindlimb | 7.31 |
| PC2 | Crown | 6.53 |
| PC2 | Dorsal Torso | 6.10 |
| PC2 | Forelimb | 4.04 |
| PC3 (eigenvalue = 1.00) | Crown | 62.03 |
| PC3 | Dorsal Torso | 21.36 |
| PC3 | Cheek | 11.47 |
| PC3 | Hindlimb | 2.63 |
| PC3 | Forelimb | 2.33 |
| PC3 | Ventral Torso | 0.18 |

Contribution to overall variance: PC1 (29.77%), PC2 (20.87%), and PC3 (16.70%). Total variance explained by PCs with eigenvalues > 1 is 67.33%.

**SI 11** PCA biplot of pelage brightness across species for the *Propithecus* genus

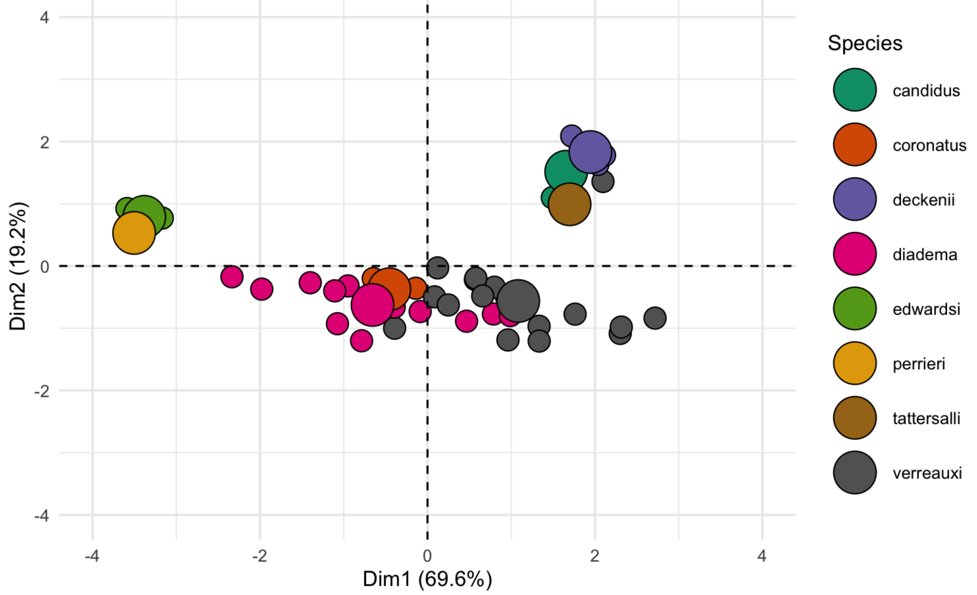

**SI 12** Loadings from the hair brightness PCA within *Propithecus* for each PC with an eigenvalue above one, from greatest to least variance explained.

| Principal Component | Variable | Variance Explained (%) |
|---|---|---|
| PC1 (eigenvalue = 1.86) | Hindlimb | 24.57 |
| PC1 | Dorsal torso | 24.21 |
| PC1 | Forelimb | 23.68 |
| PC1 | Tail | 20.23 |
| PC1 | Crown | 7.32 |

Contribution to overall variance: PC1 (64%)

**SI 13** PCA biplot of pelage hue across species for the *Propithecus* genus

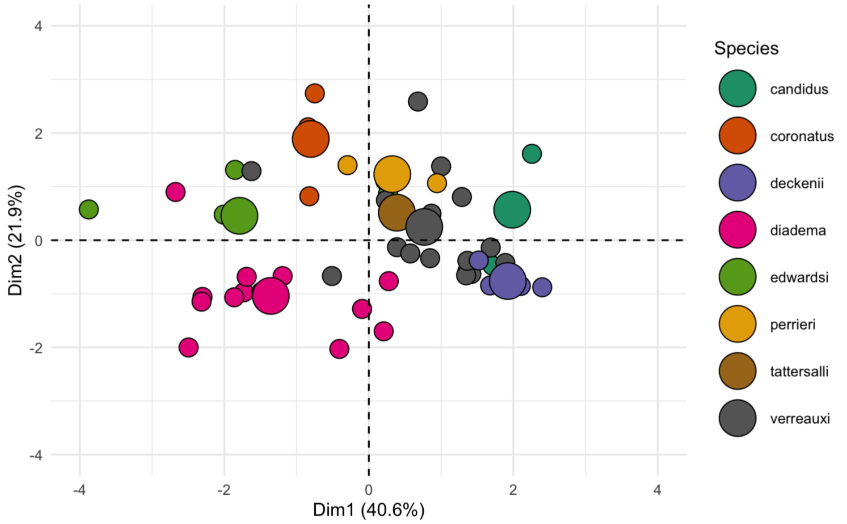

**SI 14** Loadings from the hair hue PCA within *Propithecus* for each PC with an eigenvalue above one, from greatest to least variance explained.

| Principal Component | Variable | Variance Explained (%) |
|---|---|---|
| PC1 (eigenvalue = 1.56) | Forelimb | 30.71 |
| PC1 | Dorsal torso | 29.51 |
| PC1 | Hindlimb | 26.53 |
| PC1 | Ventral torso | 8.65 |
| PC1 | Cheek | 4.48 |
| PC1 | Crown | 0.03 |
| PC2 (eigenvalue = 1.15) | Cheek | 42.57 |
| PC2 | Ventral torso | 32.88 |
| PC2 | Hindlimb | 15.39 |
| PC2 | Crown | 5.58 |
| PC2 | Forelimb | 3.56 |
| PC2 | Dorsal torso | 0.01 |
| PC3 (eigenvalue = 1.01) | Crown | 88.57 |
| PC3 | Cheek | 5.00 |
| PC3 | Dorsal torso | 3.87 |
| PC3 | Ventral torso | 2.26 |
| PC3 | Forelimb | 0.30 |
| PC3 | Hindlimb | 0.001 |

Contribution to overall variance: PC1 (40.57%), PC2 (21.90%), PC3 (17.05%), and PC4 (14.30%). Total variance explained by PCs with eigenvalues > 1 is 79.52%.

**SI 15** PCA biplot of pelage hair and follicle density across all Indriidae taxa

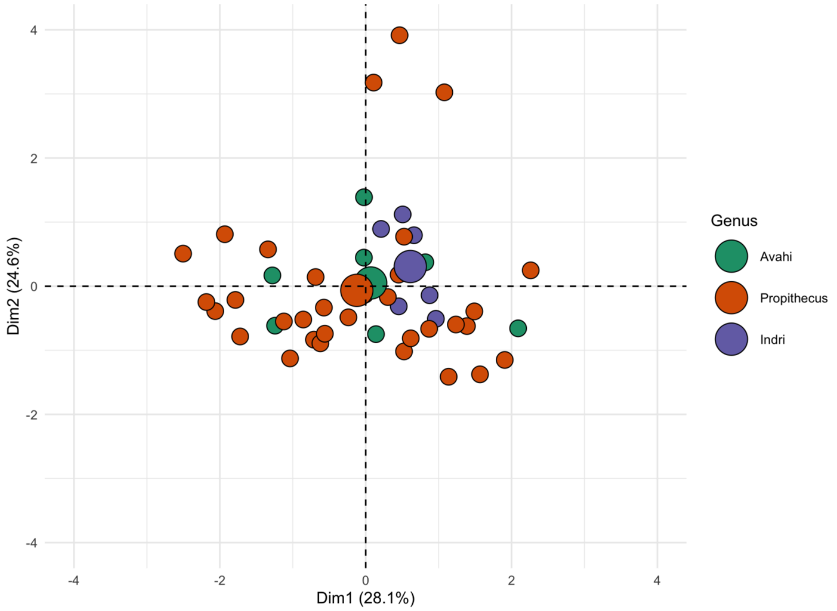

**SI 16** Loadings from the hair and follicle density PCA across all Indriidae taxa, for each PC with an eigenvalue above one, from greatest to least variance explained.

| Principal Component | Variable | Variance Explained (%) |
|---|---|---|
| PC1 (eigenvalue = 1.18) | Forelimb, follicle density | 45.13 |
| PC1 | Hindlimb, follicle density | 44.07 |
| PC1 | Crown, follicle density | 9.89 |
| PC1 | Ventral torso, hair density | 0.91 |
| PC1 | Dorsal torso, hair density | < 0.01 |
| PC2 (eigenvalue = 1.11) | Ventral torso, hair density | 56.30 |
| PC2 | Crown, follicle density | 24.34 |
| PC2 | Hindlimb, follicle density | 11.76 |
| PC2 | Forelimb, follicle density | 4.60 |
| PC2 | Dorsal torso, hair density | 2.99 |
| PC3 (eigenvalue = 1.00) | Dorsal torso, hair density | 92.31 |
| PC3 | Crown, follicle density | 6.81 |
| PC3 | Hindlimb, follicle density | 0.65 |
| PC3 | Forelimb, follicle density | 0.16 |
| PC3 | Ventral torso, hair density | 0.06 |

Contribution to overall variance: PC1 (28.07%), PC2 (24.63%), and PC3 (20.03%). Total variance explained by PCs with eigenvalues > 1 is 72.72%.

**SI 17** PCA biplot of pelage hair and follicle density within *Propithecus*

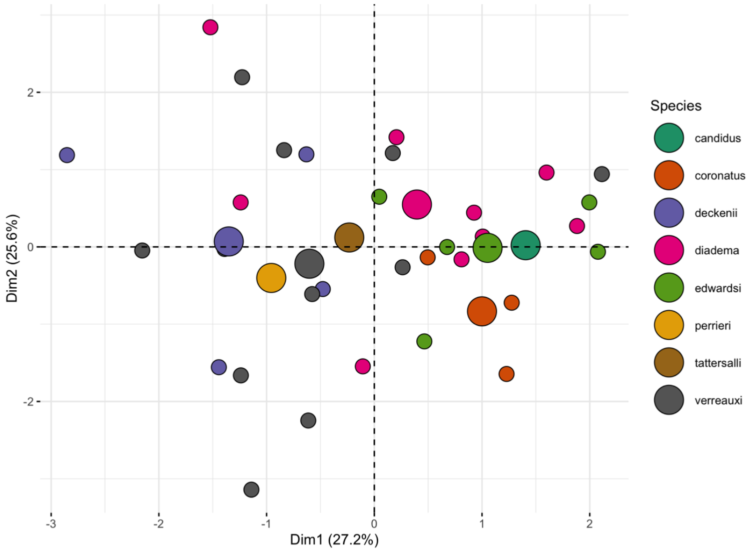

**SI 18** Loadings from the hair and follicle density PCA within *Propithecus*, for each PC with an eigenvalue above one, from greatest to least variance explained.

| Principal Component | Variable | Variance Explained (%) |
|---|---|---|
| PC1 (eigenvalue = 1.28) | Forelimb, follicle density | 30.84 |
| PC1 | Crown, hair density | 23.01 |
| PC1 | Hindlimb, follicle density | 22.54 |
| PC1 | Crown, follicle density | 18.91 |
| PC1 | Hindlimb, hair density | 3.49 |
| PC1 | Ventral torso, hair density | 1.20 |
| PC2 (eigenvalue = 1.24) | Crown, follicle density | 30.00 |
| PC2 | Hindlimb, follicle density | 21.84 |
| PC2 | Hindlimb, hair density | 20.22 |
| PC2 | Ventral torso, hair density | 14.74 |
| PC2 | Crown, hair density | 12.14 |
| PC2 | Forelimb, follicle density | 1.07 |
| PC3 (eigenvalue = 1.03) | Ventral torso, hair density | 61.36 |
| PC3 | Crown, hair density | 17.01 |
| PC3 | Forelimb, follicle density | 11.80 |
| PC3 | Hindlimb, hair density | 9.48 |
| PC3 | Crown, follicle density | 0.28 |
| PC3 | Hindlimb, follicle density | 0.06 |

Contribution to overall variance: PC1 (27.24%), PC2 (25.56%), and PC3 (17.68%). Total variance explained by PCs with eigenvalues > 1 is 70.48%.

**SI 19** Summary of ANOVA examining hair and follicle density variation across distinct body regions for Propithecus.

| | DF | Sum Sq. | Mean Sq. | F value | P value |
|---|---|---|---|---|---|
| *Hair density* | | | | | |
| Body Region | 4 | 605041 | 151260 | 3.968 | **0.009** |
| Residuals | 35 | 1334233 | 38121 | | |
| *Follicle density* | | | | | |
| Body Region | 4 | 82046 | 20511 | 1.357 | 0.269 |
| Residuals | 35 | 529130 | 15118 | | |

**SI 20** Boxplots illustrating variation in (A) hair density and (B) follicle density across body regions, for *Propithecus*. Black lines represent the average, and black dots represent outliers.

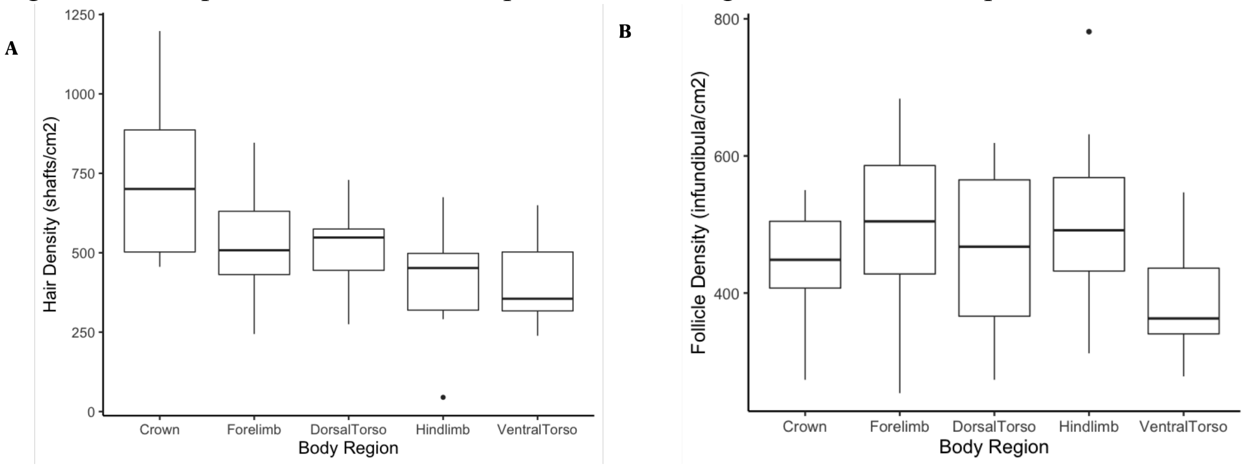


**SI 21** PCA biplot of pelage hue across the difference in spectral sensitivity in each sampled population, for diurnal Indriids (*Propithecus* and *Indri*)

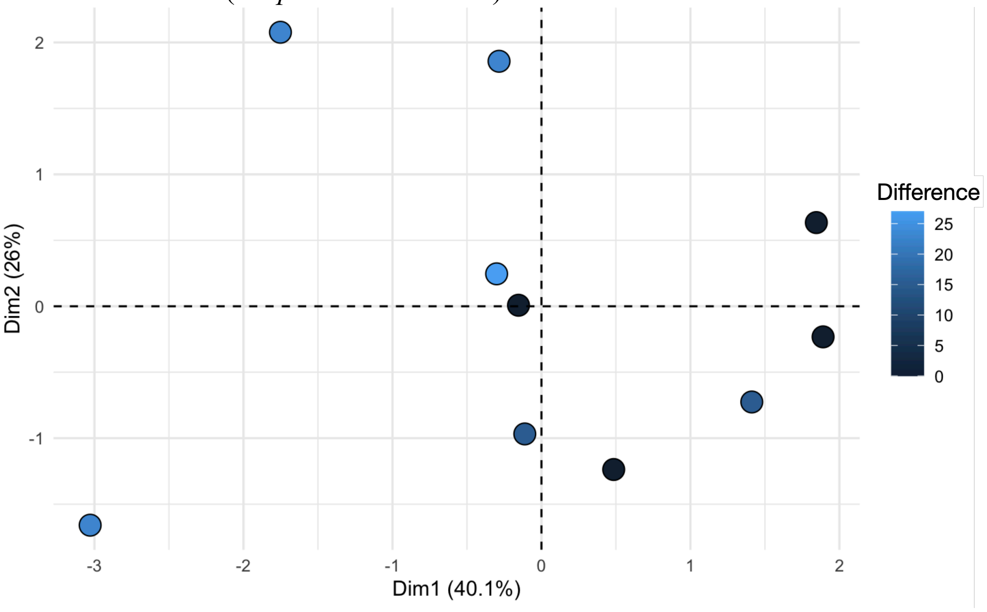

**SI 22** Loadings from the hair hue PCA used for the opsin PGLMM within Indriidae, for each PC with an eigenvalue above one, from greatest to least variance explained.

| Principal Component | Variable | Variance Explained (%) |
|---|---|---|
| PC1 (eigenvalue = 1.55) | Forelimb | 36.71 |
| PC1 | Hindlimb | 31.77 |
| PC1 | Dorsal torso | 21.82 |
| PC1 | Cheek | 6.60 |
| PC1 | Crown | 3.04 |
| PC1 | Ventral torso | 0.05 |
| PC2 (eigenvalue = 1.25) | Ventral torso | 42.18 |
| PC2 | Cheek | 35.44 |
| PC2 | Hindlimb | 6.58 |
| PC2 | Dorsal torso | 5.35 |
| PC2 | Crown | 5.19 |
| PC2 | Forelimb | 2.26 |

Contribution to overall variance: PC1 (40.11%), and PC2 (25.98%). Total variance explained by PCs with eigenvalues > 1 is 66.09%.

**SI 23** Results of PGLMMs predicting pelage hue across diurnal Indriids, against the difference in spectral sensitivity and total number of opsin alleles in the populations.

| Variable | Mean | Lower CI | Upper CI | ESS | P MCMC |
|---|---|---|---|---|---|
| PC1; $H^2 = 0.409$ | | | | | |
| Intercept | 1.584 | 0.629 | 2.553 | 9,000 | 0.011 |
| Difference | -0.110 | -0.250 | 0.023 | 9,000 | 0.095[!] |
| NoOfAlleles | 2.364 | -1.364 | 5.816 | 8,664 | 0.165 |
| PC2; $H^2 = 0.735$ | | | | | |
| Intercept | 0.625 | -2.600 | 3.975 | 8,471 | 0.567 |
| Difference | 0.085 | -0.253 | 0.448 | 9,000 | 0.572 |
| NoOfAlleles | -2.464 | -10.774 | 5.466 | 9,942 | 0.506 |

Bold values indicate significance below 0.05, [!] indicates $p < 0.10$, $H^2$ is akin to phylogenetic signal.

**SI 24** A map of Madagascar indicating the interaction between the temperature and rainfall gradients, as derived from previous studies (Irwin, 2006; Kamilar & Muldoon, 2010).

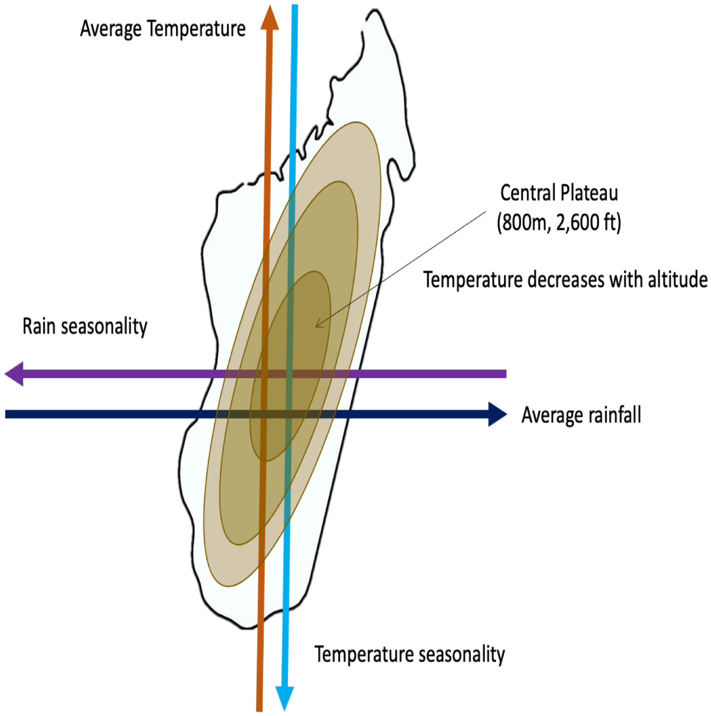


**References to supplementary figures and tables**

Irwin, M. T. (2006). Ecologically enigmatic lemurs: The sifakas of the Eastern foress (Propithecus candidus, P. diadema, P. edwardsi, P. perrieri, and P. tattersalli). In *Lemurs: Ecology and Adaptation* (pp. 305–326).

Kamilar, J. M., & Muldoon, K. M. (2010). The climatic niche diversity of Malagasy primates: A phylogenetic perspective. *PLoS ONE*, *5*(6). https://doi.org/10.1371/journal.pone.0011073
